# A malaria parasite phospholipase facilitates efficient asexual blood stage egress

**DOI:** 10.1101/2023.03.13.532312

**Authors:** Abhinay Ramaprasad, Paul-Christian Burda, Konstantinos Koussis, James A Thomas, Emma Pietsch, Enrica Calvani, Steven A Howell, James I MacRae, Ambrosius P. Snijders, Tim-Wolf Gilberger, Michael J Blackman

## Abstract

Malaria parasite release (egress) from host red blood cells involves parasite-mediated membrane poration and rupture, thought to involve membrane-lytic effector molecules such as perforin-like proteins and/or phospholipases. With the aim of identifying these effectors, we disrupted the expression of two *Plasmodium falciparum* perforin-like proteins simultaneously and showed that they have no essential roles during blood stage egress. Proteomic profiling of parasite proteins discharged into the parasitophorous vacuole (PV) just prior to egress detected the presence in the PV of a lecithin:cholesterol acyltransferase (LCAT; PF3D7_0629300). Conditional ablation of LCAT resulted in abnormal egress and a reduced replication rate. Lipidomic profiles showed drastic changes in several phosphatidylserine and acylphosphatidylglycerol species during egress. We thus show that, in addition to its previously demonstrated role in liver stage merozoite egress, LCAT is required to facilitate efficient egress in asexual blood stage malaria parasites.

**Author Summary:** Malaria kills over half a million people every year worldwide. It is caused by a single-celled parasite called *Plasmodium falciparum* that grows and multiplies within a bounding vacuole, inside red blood cells of the infected individuals. Following each round of multiplication, the infected cell is ruptured in a process known as egress to release a new generation of parasites. Egress is required for the disease to progress and is orchestrated by the parasite. The parasite sends out various molecules to puncture and destroy the membranes of the vacuole and the red blood cell. However, little is known about these molecules. In this work, we set out to identify these molecules by using genetic and proteomics approaches. We screened the molecules the parasite sends out during egress and identified a parasite enzyme called LCAT present in the vacuole. Our experiments found that mutant parasites that were unable to make LCAT clumped together and could not escape the infected cell properly. As a result, we saw a reduction in the rate at which these parasites spread through the red blood cells. Taken together, our findings suggest that *P. falciparum* needs LCAT to efficiently break out of red blood cells.

## Introduction

During the asexual blood stages (ABS) of their life cycle, malaria parasites grow and replicate asexually within a parasitophorous vacuole (PV) in host red blood cells (RBCs). At the end of each cycle of intraerythrocytic development, invasive merozoites are released from the host cell in a coordinated lytic process known as egress, to invade fresh RBCs. Egress involves a rapid sequence of events resulting in the rupture of the two bounding membranes, the PV membrane (PVM) and the RBC membrane (RBCM). Minutes before egress, the PV rounds up without swelling in a calcium-dependent process (Glushakova et al., 2018), followed by PVM leakage and rupture, then subsequent poration and rupture of the RBCM (Abkarian et al., 2011; Glushakova *et al*., 2018; Glushakova et al., 2010; Glushakova et al., 2005; Hale et al., 2017; Wickham et al., 2003). Egress is initiated by activation of a cGMP-dependent protein kinase (PKG) (Collins et al., 2013b) in coordination with a calcium-dependent protein kinase called CDPK5 (Absalon et al., 2018) by triggering the discharge of specialised parasite secretory organelles called micronemes and exonemes. A subtilisin-like parasite protease (SUB1) is released from the exonemes into the PV lumen where it initiates a proteolytic cascade by cleaving and activating several effector molecules including members of the papain-like SERA protein family and several components of the merozoite surface (Das et al., 2015; Koussis et al., 2009; Ruecker et al., 2012; Silmon de Monerri et al., 2011; Yeoh et al., 2007). Activated SERA6 precisely cleaves β-spectrin in the RBC cytoskeleton to bring about the final step of RBCM rupture (Thomas et al., 2018). Despite these many insights, the molecular basis for the events that precede RBCM rupture, PVM rupture and RBCM poration, remains unknown. Membrane-lytic effector molecules such as perforin-like proteins (PLPs) and phospholipases likely bring about these membrane-degradative events. Previous evidence (Collins et al., 2017) showing that RBCM poration occurs immediately following PVM rupture has led us to postulate that the same effector molecules may mediate both events. These effectors may either be released from secretory organelles just prior to egress or are constitutively resident within the PV waiting to be activated by the SUB1-initiated proteolytic cascade.

Pore-forming proteins of the membrane attack complex component/perforin (MACPF) superfamily disrupt membranes by inserting their characteristic MACPF domain into the target phospholipid bilayer to form a transmembrane channel (Dal Peraro and van der Goot, 2016). Five *Plasmodium* proteins possessing MACPF domains, annotated as perforin-like proteins (PPLPs), are expressed significantly in the sexual gametocyte stages that transmit the parasite to the mosquito vector, as well as in the mosquito ookinete and sporozoite stages; however, the PPLPs are transcribed only at low levels in ABS (PlasmoDB v46, (Kaiser et al., 2004)). Consistent with this, in previous genetic analyses in which each of the five PPLP genes was individually disrupted, the genes were shown to be dispensable for ABS growth but were instead implicated in other parasite life cycle transitions (Guerra and Carruthers, 2017). PPLP1 is required for sporozoite egress from transient vacuoles formed by the parasite in host hepatocytes (Ishino et al., 2005; Risco-Castillo et al., 2015; Yang et al., 2017), while PPLP2 is required for gametocyte egress (Deligianni et al., 2013; Wirth et al., 2014). Despite these findings, other studies have shown that PPLP1 and PPLP2 are detectable in ABS parasites (Garg et al., 2013; Wirth *et al*., 2014) and biochemical and small molecule inhibitor evidence suggested a possible role for both PPLPs in membrane poration during merozoite egress (Garg *et al*., 2013; Garg et al., 2020). One plausible explanation for this apparent discrepancy is that PPLP1 and PPLP2 perform redundant functions in ABS, in which case disruption of any one gene might be functionally complemented by the other. However, this possibility has not been examined.

Phospholipases (PLs) mediate membrane lysis through hydrolytic cleavage of either of the acyl chains (phospholipases A1, A2 and B) or phosphodiester bonds in the glycerol backbone (phospholipases C and D) of membrane phospholipids. Phospholipases often aid exit of intracellular bacteria from the cellular vacuole (Hybiske and Stephens, 2008), and can also cleave fatty acids from phospholipids to alter membrane curvature through localised phospholipid asymmetry (Zimmerberg and Kozlov, 2006), raising the possibility of similar roles during the extensive morphological changes associated with malarial egress. A systematic functional analysis of 20 out of 27 putative phospholipases in the human malaria parasite, *Plasmodium falciparum* suggested a high degree of functional redundancy, with only five being found to possibly be essential in ABS (Burda et al., 2021a). Of those phospholipases deemed dispensable in ABS, a secreted phospholipase with a lecithin:cholesterol acyltransferase (LCAT)-like domain has previously been shown to disrupt membranes during mosquito and liver stages of the parasite in the rodent malarial species *P. berghei* (Bhanot et al. 2005; Burda et al. 2015). LCAT was found to be expressed on the surface of sporozoites, and LCAT-null sporozoites had a reduced capacity to egress from oocysts and migrate through host hepatocytes. LCAT localizes to the PV and PVM following invasion of hepatocytes and LCAT-null parasites were defective in liver stage schizont egress due to impaired PVM rupture that prevents or delays merozoite release. Whether LCAT plays any role in *Plasmodium* blood stage egress remains unclear.

Here, we first rule out any essential requirement for PPLP1 and PPLP2 in ABS egress. We then profile the repertoire of proteins that are discharged from the micronemes and exonemes at egress to examine whether they include any previously unidentified effector molecule(s) potentially involved in PVM rupture and RBC poration. Our resulting work allows us to demonstrate that LCAT plays a previously unrecognised role in facilitating efficient egress.

## Results

### Perforin-like proteins are dispensable for ABS egress

While genetic ablation of PPLP1 and PPLP2 individually has previously been shown to have no effect on parasite proliferation (Wirth *et al*., 2014; Yang *et al*., 2017), the question remained as to whether the proteins could function together to mediate membrane poration during egress. To examine this, we attempted to generate a parasite line in which both PPLP1 (PF3D7_0408700) and PPLP2 (PF3D7_1216700) were simultaneously disrupted. To do this, we set about floxing the MACPF-encoding domains of both genes in the DiCre-expressing *P. falciparum* line B11 (Figure 1A and Supplementary Figure S1A), so that rapamycin (RAP)-mediated activation of the DiCre could be used to simultaneously excise functionally critical regions of both genes. Due to an error in repair plasmid design for *pplp1*, we inadvertently introduced a frameshift mutation and disrupted the gene when floxing its MACPF domain. The resultant PPLP1-null *P. falciparum* line (D1) displayed normal growth (Supplementary Figure S1B) as expected, and so was used as the background for subsequent floxing of *pplp2*. RAP treatment of the final modified parasite line (called PPLP1:loxNint/PPLP2:loxPint) led to efficient excision of both genetic loci within a single erythrocytic cycle (Figure 1B). As shown in (Figure 1C, D), the resulting PPLP1/PPLP2-null mutant parasites displayed normal proliferation rates with no discernible effect on parasite development. Importantly, the PPLP1/PPLP2-null schizonts also underwent typical RBC poration just prior to egress (Supplementary Movie S1). It was concluded unambiguously that neither PPLP1 nor PPLP2 alone or in combination have an essential role in membrane poration or RBC rupture during ABS egress.

**Figure 1.**
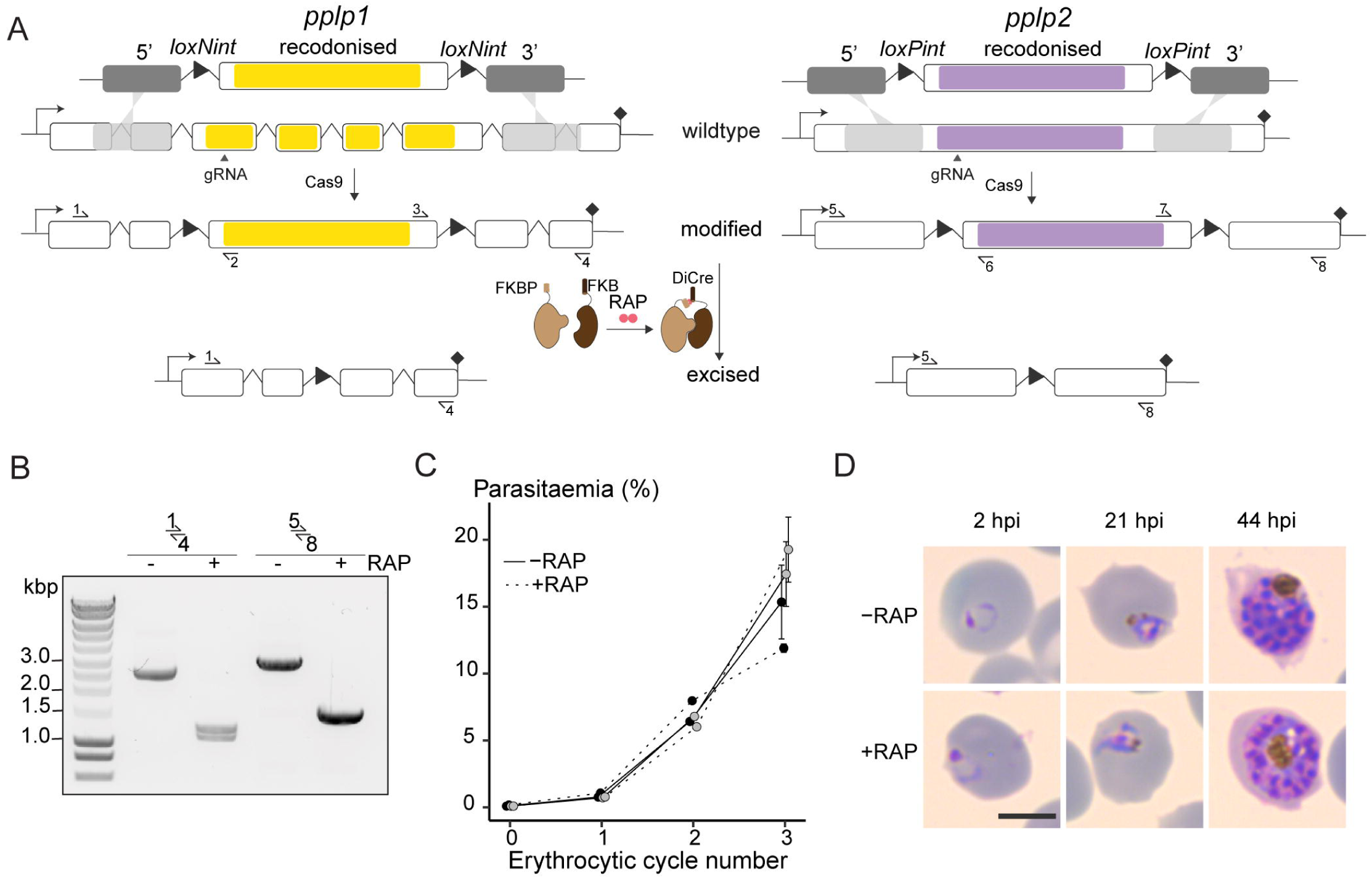
*P. falciparum* PPLP1 and PPLP2 are dispensable for asexual blood stage egress. **A)** Strategy used for simultaneous conditional disruption of both PPLP1 and PPLP2 in the parasite line PPLP1:loxNint/PPLP2:loxPint. The MACPF domains of both PLPs (yellow and purple) are floxed by introducing loxN- and loxP-containing introns (loxPints) respectively. Site of targeted Cas9-mediated double-stranded DNA break (marked “gRNA”), left and right homology arms for homology-directed repair (5’ and 3’) and diagnostic PCR primers (half-arrows 1 to 8) are indicated. RAP-induced dimerization of N- and C-terminal subunits of the Cre enzyme enables Cre-mediated excision of the floxed regions rendering both the genes non-functional. **B)** Diagnostic PCR using primers 1-4 and 5-8 (representative of 3 independent experiments) confirms efficient excision at both loci sampled at 12 hours post-RAP (+RAP) or -mock treatment (-RAP) of ring stages. **C)** Replication of mock-(solid line) and RAP-treated (dashed line) parasites from two clonal lines of PPLP1:loxNint/PPLP2:loxPint, C1 and C2, over three erythrocytic cycles (error bars, ± SD, triplicate RAP treatments with different blood sources). There is no significant difference in replication rates. **D)** Light microscopic images of Giemsa-stained PPLP1:loxNint/PPLP2:loxPint parasites following mock- or RAP-treatment at ring stage in cycle 0 (representative of 2 independent experiments). PPLP1/PPLP2-null parasites exhibit normal parasite development. Scale bar, 5 µm.

### Differential proteomics of SUB1-null schizonts identifies proteins discharged into the PV at egress

Minutes before merozoite egress, malaria parasites discharge the contents of micronemes and exonemes onto the merozoite surface or into the PV lumen, where some of the discharged proteins perform specific tasks to bring about egress. Discharge of the exoneme protease SUB1 is required to mediate both PVM and RBCM rupture (Thomas *et al*., 2018). As a consequence, SUB1-null parasites arrest in a state in which microneme and exoneme discharge has occurred normally, but the PVM and RBCM remain intact, resulting in the discharged organelle contents remaining trapped within an intact PV. This is in contrast to treatment of schizonts with the PKG inhibitor 4-[7-[(dimethylamino)methyl]-2-(4-fluorphenyl)imidazo[1,2-α]pyridine-3-yl]pyrimidin-2 amine (compound 2, C2); this leads to a similar morphological phenotype, with merozoites trapped within an intact PVM and RBCM, but in this case microneme/exoneme discharge is blocked (Figure 2A). We reasoned that these contrasting phenotypes provided an opportunity to selectively identify the repertoire of microneme and exoneme proteins that are discharged into the PV at egress. Of particular interest, we reasoned that this group of discharged proteins might include amongst them previously unidentified effector molecule(s) involved in PVM rupture and RBC poration.

**Figure 2.**
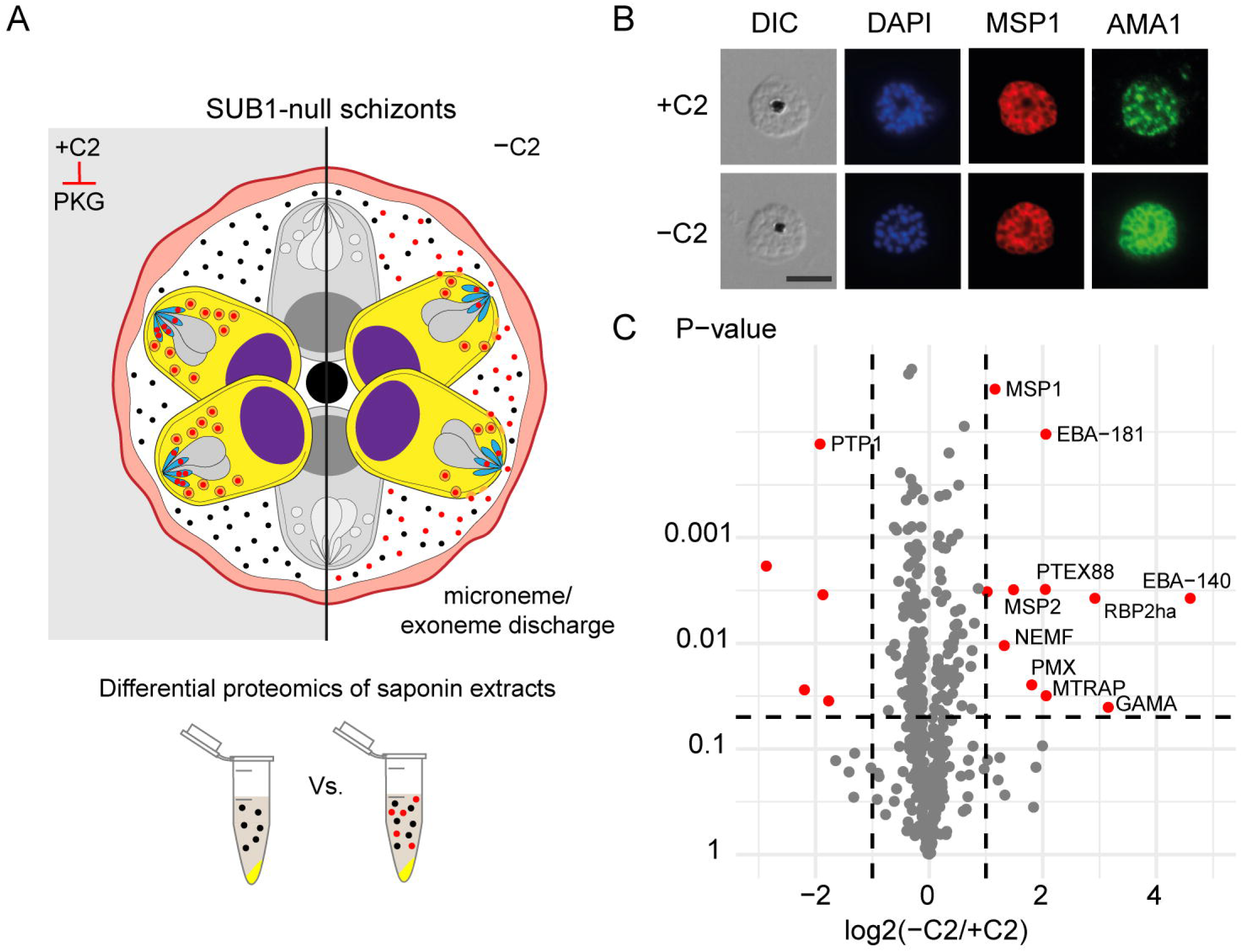
Proteomic identification of organelle proteins discharged during egress. **A)** Schematic of experimental design to identify components of micronemes and exonemes released prior to egress. SUB1-null schizonts were incubated in C2 (+C2) to prevent discharge of microneme and exoneme proteins (marked in red). Upon washing off C2 (-C2), micronemes and exonemes release their contents into the PV and are retained there in SUB1-null schizonts. Comparisons of the proteomic profiles of saponin extracts of the C2-arrested and C2-washed SUB1-null schizonts is expected to show differences in the abundance of microneme and exoneme proteins. **B)** Immunofluorescence assay showing AMA1 localization (green). In the presence of C2, AMA1 is restricted to the micronemes at the apical ends of the merozoites. In contrast, upon C2 removal, AMA1 translocates onto the merozoite cell surface. Parasite surface marker MSP1 (red), DAPI-stained nuclei (blue) and DIC, differential interference contrast are also shown. Scale bar, 5 µm. **C)** Volcano plot showing enrichment of 12 proteins (red) in the C2-washed schizonts compared to C2-arrested SUB1-null schizonts (values averaged from biological triplicates).

To selectively identify these ‘trapped’ components, we allowed SUB1-null schizonts to mature in the presence of C2, then washed away the C2 for 20 minutes to allow organelle discharge before finally releasing the PV contents using saponin lysis, which disrupts both the RBCM and PVM but not the parasite plasma membrane. Successful microneme discharge upon washing away C2 was confirmed by observing relocalisation of the micronemal protein AMA1 to the periphery of merozoites (Figure 2B). We then compared the proteomic profiles of the saponin lysates of C2-washed (-C2) and C2-arrested (+C2) parasites to identify those exonemal/micronemal proteins discharged into the PV within the 20-minute window.

A total of 503 parasite proteins were identified, including several established constitutively PV-resident proteins like the SERA family of papain-like proteins (Miller et al., 2002; Ruecker *et al*., 2012) and protein phosphatase UIS2 (Khosh-Naucke et al., 2018), and proteins exported to the RBC cytosol including secreted rhoptry proteins and the PHIST family of exported proteins (Supplementary Table S2). The remaining proteins were known parasite cytosolic or nuclear proteins including ribosomal and proteasomal proteins, probably representing contaminants from the parasite fraction. Levels of all these proteins were largely equivalent between the +C2 and -C2 schizonts, as expected for constitutively-expressed PV proteins. A total of only 12 proteins were found to be significantly more abundant in -C2 saponin extracts (p<0.05, more than 2-fold change; see Figure 2B). These included the exoneme protease plasmepsin X (PMX) (Nasamu et al., 2017; Pino et al., 2017) which was enriched 4-fold, as well as four membrane proteins that are discharged from micronemes during schizont rupture: EBA-140, EBA-181, GAMA and MTRAP (4-fold to 24-fold enrichment). The enrichment of several bona fide microneme/exoneme proteins in the -C2 samples indicated that our strategy was successful, but no candidates with established membranolytic activity was identified amongst the enriched (organelle-derived) population. However, a lecithin:cholesterol acyltransferase (LCAT) previously implicated in liver stage merozoite egress in the rodent malaria parasite *Plasmodium berghei* (Burda et al., 2015), was detected in all the samples (Supplementary Table S2) indicating that this is constitutively resident in the PV. We chose this protein for further investigation.

### Conditional genetic ablation of LCAT reduces blood stage proliferation

Previous transcriptomic analyses indicate that peak expression of LCAT (PF3D7_0629300) during ABS occurs during schizont development (Lopez-Barragan et al., 2011). To confirm the localisation of LCAT in ABS *P. falciparum* parasites, we appended a C-terminal GFP-tag to the endogenous gene using the selection-linked integration (SLI) system (Birnbaum et al., 2017) (Figures 3A and Supplementary Figure S2A and B). Live fluorescence microscopy of LCAT-GFP schizonts revealed that the fluorescent signal localized to focal structures within the parasite and around developing merozoites, pointing towards a localization to secretory organelles and the PV (Figure 3A). A similar localization was observed by immunofluorescence analysis (IFA) of fixed parasites expressing LCAT fused to spaghetti monster-Myc (smMyc) tag (Viswanathan et al., 2015) (Supplementary Figure S2C, D and E). IFA using rabbit polyclonal antibodies raised against LCAT similarly confirmed its localization to secretory organelles and to the PV (indicated by partial co-localization with the PV protein SERA5) (Figure 3B).

**Figure 3.**
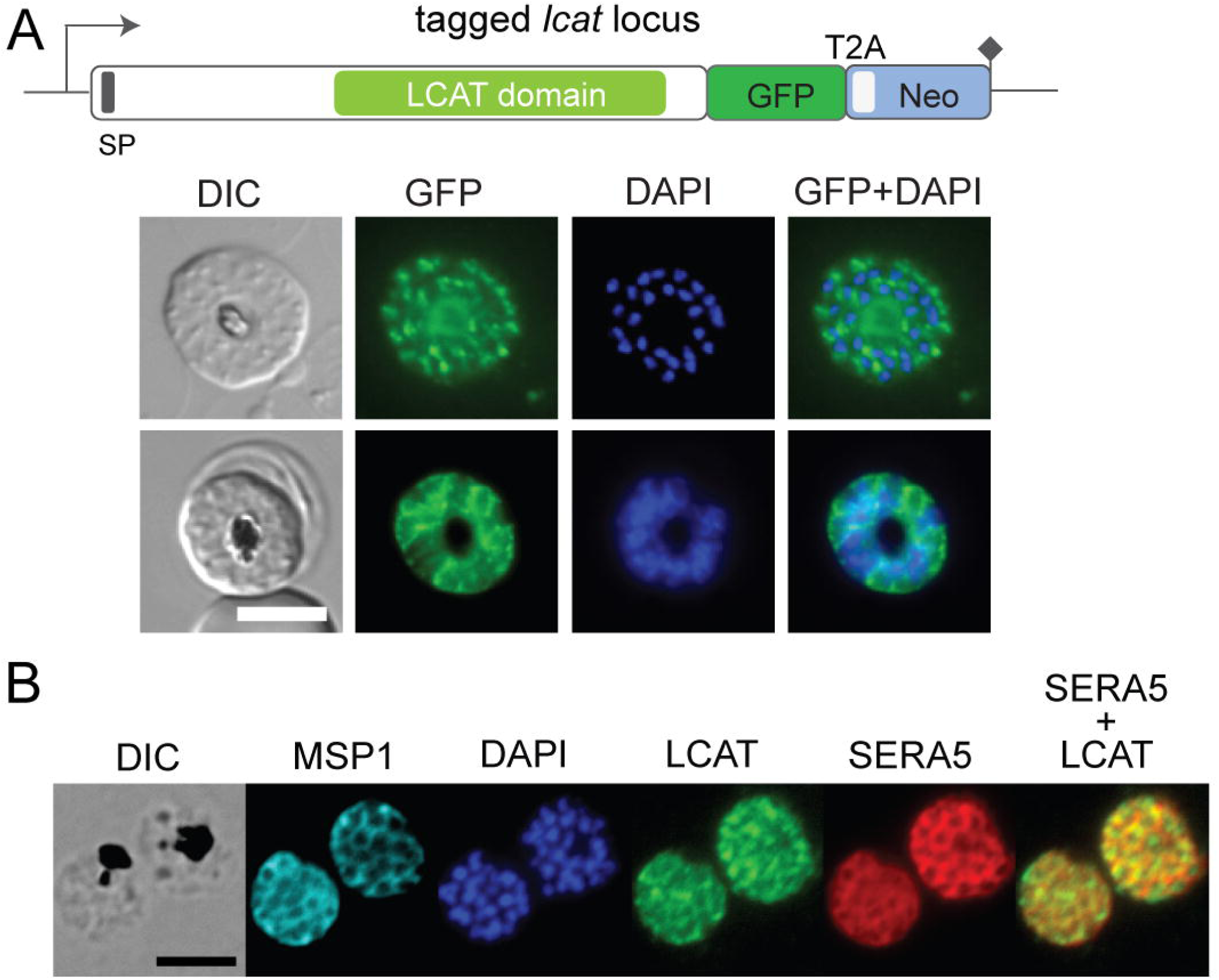
LCAT localises to secretory vesicles and the PV in blood stage schizonts. **A)** Live-cell microscopy of late schizonts expressing endogenously tagged LCAT-GFP (green). Nuclei were stained with DAPI (blue). DIC, differential interference contrast. Scale bar, 5 µm. **B)** IFA using anti-LCAT polyclonal antibodies showing that LCAT partially colocalises with the PV protein SERA5. Scale bar, 5 µm.

To conditionally ablate the *lcat* gene, we designed a DiCre-mediated gene disruption strategy that introduces a translational frameshift, truncating the gene to render it non-functional (Figure 4A and Supplementary Figure S3). For this, we floxed a short 200 bp segment upstream of the putative catalytic domain in the *lcat* gene (PF3D7_0629300) by introducing two closely-opposed loxPint modules, producing a modified inducible LCAT knockout line called LCAT:2loxPint. DiCre-mediated excision of the floxed region was predicted to result in a frameshift mutation that introduces multiple stop codons in the downstream sequence encoding the LCAT catalytic domain. RAP-treatment of two clonal LCAT:2loxPint lines (B10 and F10) resulted in efficient excision of the floxed sequence (Figure 4B) and loss of LCAT expression as confirmed by western blotting and IFA (Figure 4C and D).

**Figure 4.**
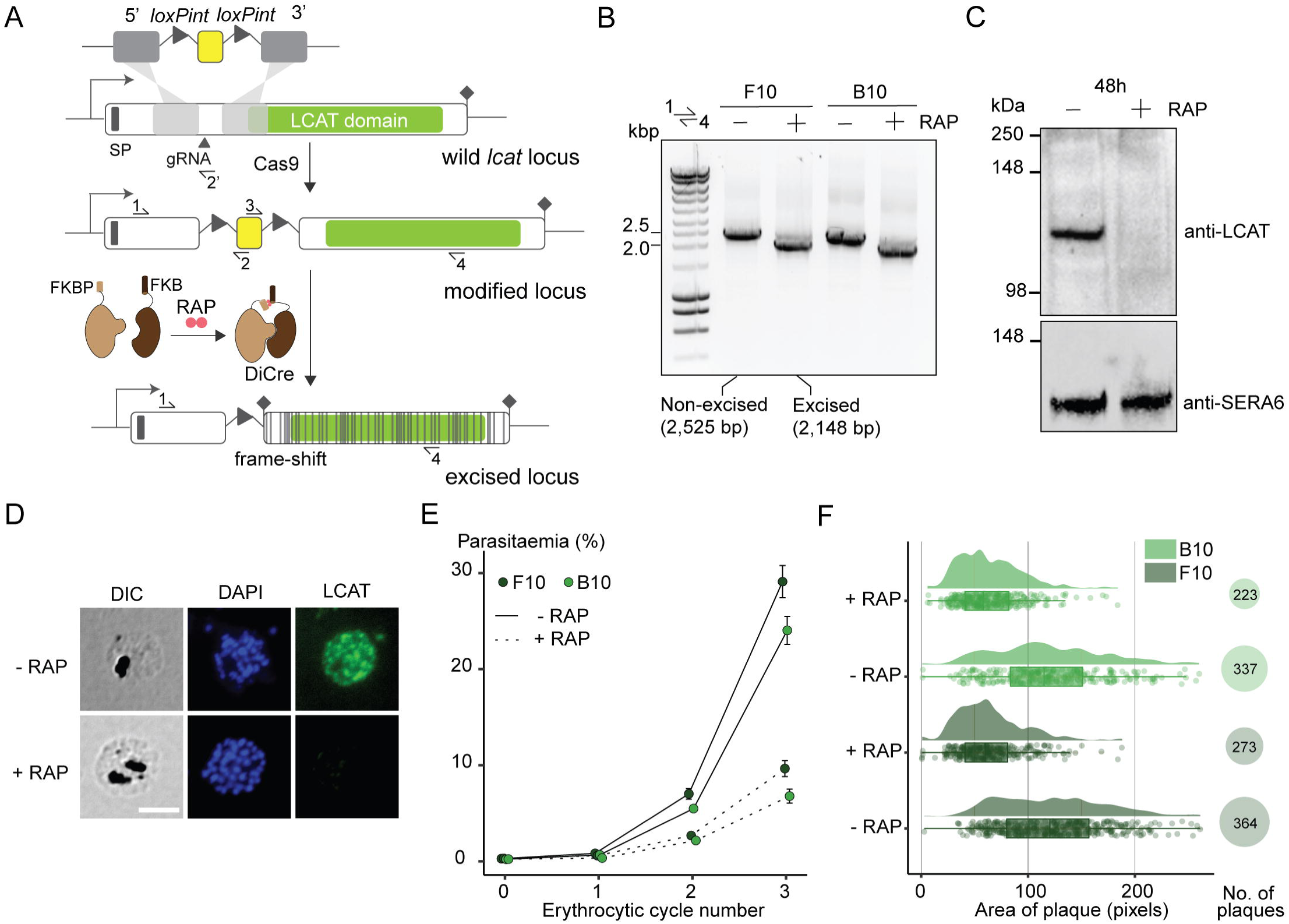
Genetic ablation of LCAT expression reduces blood stage proliferation. **A)** Strategy used for conditional disruption of LCAT in parasite line LCAT:2loxPint. A 200bp region upstream of the predicted LCAT domain (green) is floxed by introducing two loxPint modules. Predicted secretory signal peptide (SP), site of targeted Cas9-mediated double-stranded DNA break (marked “gRNA”), left and right homology arms for homology-directed repair (5’ and 3’), recodonized sequence (yellow) and diagnostic PCR primers (half arrows 1-4) are indicated. RAP-induced DiCre-mediated excision results in frameshift that renders the gene non-functional (grey lines). **B)** Diagnostic PCR 12 h following mock- or RAP-treatment of ring-stage LCAT:2loxPint (representative of 3 independent experiments) confirms efficient gene excision. Expected amplicon sizes are indicated. **C)** Western blots (representative of 3 independent experiments) showing successful RAP-induced ablation of LCAT expression in LCAT:2loxPint parasites sampled at 48 h post invasion. SERA6 was probed as loading control. **D)** IFA of RAP-treated (+RAP) and mock-treated (-RAP) mature LCAT:2loxPint schizonts showing that expression of LCAT is lost following RAP treatment. Scale bar, 5 µm. **E)** RAP-treatment results in reduced replication rate in two clonal lines, F10 and B10, of LCAT:2loxPint parasites. Data shown are averages from triplicate biological replicates using different blood sources (error bars, ± SD). **F)** RAP-treatment results in reduction in both number and area of clonal plaques formed over five erythrocytic cycles (10 days of growth) in LCAT:2loxPint clonal lines (individual points represent each plaque, density plot shows distribution of these points and boxplot provides median summary statistics).

The RAP-treated LCAT-null clonal lines displayed a reduced proliferation rate (∼66% reduction over 3 erythrocytic cycles) compared to mock-treated controls (Figure 4E). Longer-term viability of the LCAT-null parasites over ∼5 erythrocytic cycles as assessed by plaque assay (Thomas et al., 2016) reflected this, with a 25-34% reduction in the number of plaques formed following RAP-treatment, as well as a significant reduction in the average area of these plaques (Figure 4F). It was concluded that LCAT is important for ABS parasite replication *in vitro*.

### Loss of LCAT causes inefficient egress

To explore in more detail the growth defect in LCAT-null parasites, we monitored their development over the course of a single erythrocytic cycle. As shown in Figure 5A, this revealed no obvious impact on intracellular growth or morphology that might explain the proliferation defect (Figure 5A). We then visualised the behaviour of RAP- and mock-treated LCAT:2loxPint schizonts as they underwent egress using time-lapse video microscopy. The mock-treated schizonts underwent the typical morphological changes associated with egress, including PVM swelling and rounding up, followed by PVM rupture within seconds, and finally RBCM rupture and merozoite release. In contrast, LCAT-null schizonts showed clear delays between these sequential events, with reduced merozoite release upon RBCM rupture, the remaining merozoites often appearing clumped together (Figure 5B-D, Supplementary Movie S2).

**Figure 5.**
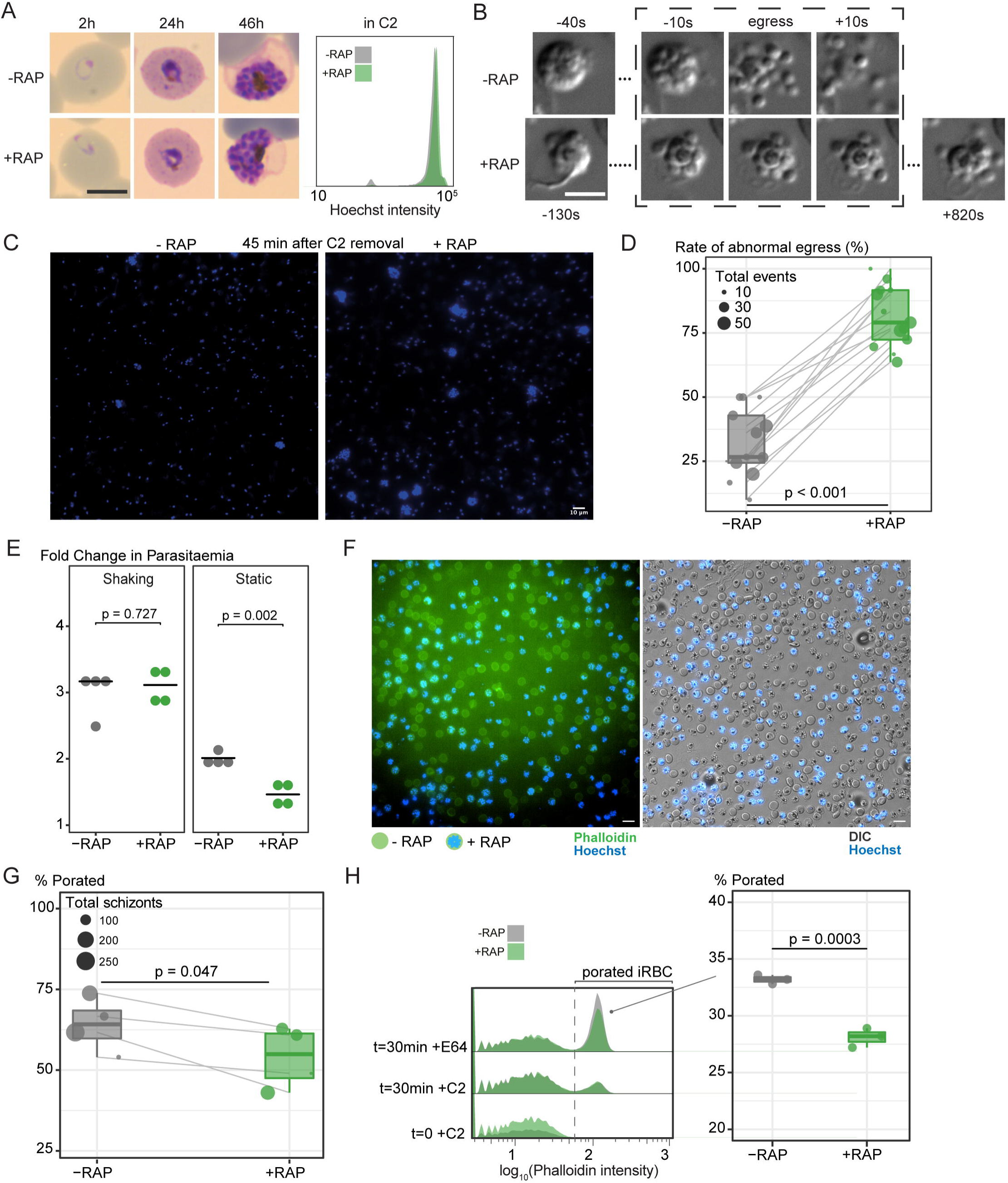
LCAT is required for efficient asexual blood stage egress. **A)** Light microscopic images of Giemsa-stained LCAT:2loxPint parasites following mock- or RAP-treatment at ring stages (representative of 2 independent experiments). LCAT-null parasites exhibit normal parasite development. Inset shows confirmation of normal growth by measuring DNA content of egress-arrested schizonts using flow cytometry. Scale bar, 5 µm. **B)** LCAT-null parasites (+RAP) show delayed onset of egress and inefficient dispersal of merozoites compared to mock-treated controls. This was defined as an abnormal egress event. Scale bar, 5 µm. **C)** DAPI-staining of egressed RAP-treated (+RAP) and mock-treated (-RAP) mature LCAT:2loxPint parasites show persistence of clumped merozoites that are products of abnormal egress in LCAT-null parasites. Scale bar, 10 µm. **D)** LCAT-null schizonts show a higher number of abnormal egress events compared to mock-treated schizonts (paired Student’s t-test). Each paired datapoint represents a 30-40 min video of RAP- and mock-treated LCAT:2loxPint schizonts (one group randomly stained with Hoechst DNA stain in each video) undergoing egress (from a total of 7 independent experiments). Size of each datapoint represents the total number of egress events (abnormal + normal) counted in the video. **E)** Fold change in parasitaemia after 4 h invasion of mock-(-RAP) and RAP-treated (+RAP) LCAT:2loxPint schizonts under shaking and static conditions. Static cultures show a significantly lower fold change in parasitaemia in RAP-treated parasites compared to mock-treated controls, while show no significant difference between the groups was observed in shaking cultures (error bars, ± SD, four replicate RAP treatments with different blood sources; individual points represent each replicate). **F)** Poration of the RBCM occurred in both mock-(RAP-) and RAP-treated (+RAP; stained blue with Hoechst DNA stain) LCAT:2loxPint schizonts as visualized using phalloidin (green) in the presence of E64 that inhibits the final step of RBCM rupture. Scale bar, 10 µm. **G)** A subtle decrease in the rate of RBCM poration (paired Student’s t-test) was observed in RAP-treated schizonts compared to mock-treated schizonts. Each paired datapoint represents a 30-40 min video of RAP- and mock-treated LCAT:2loxPint schizonts (one group randomly stained with Hoechst DNA stain in each video) undergoing egress in the presence of E64 (from a total of 3 independent experiments). Size of each datapoint represents the total number of schizonts counted in the video. **H)** Flow cytometry analysis showed emergence of porated parasitized RBCs (iRBCs) that emit higher fluorescence intensity from phalloidin following 30 min of egress in the presence of E64. A slight but consistent decrease in proportion of porated (iRBCs) was observed in RAP-treated compared to mock-treated schizonts.

To determine whether the egress phenotype contributes to the lower replication rates in the LCAT-null parasites, fold changes in parasitaemia during egress and invasion were compared in RAP- and mock-treated cultures (Figure 5E). In standard static cultures, RAP-treated parasites showed a two-fold reduction in parasitaemia increase compared to control cultures. This defect was rescued when the experiment was performed under shaking conditions, suggesting that the clumped LCAT-null merozoites can be released and dispersed efficiently under conditions of shear stress. These results also indicated that LCAT-null parasites do not have an intrinsic invasion defect. Taken together, our results show that LCAT-null parasites display a reduced replication rate that can be solely ascribed to inefficient egress from the host RBC.

To determine whether the abnormal egress phenotype of LCAT-null parasites was due to a defect in membrane poration, we visualized RBCM poration using fluorescent phalloidin, a cyclic peptide that binds to the short F-actin filaments within the RBC cytoskeleton upon poration. To facilitate visualisation of this, the experiments were performed in the additional presence of the cysteine protease inhibitor E64 which blocks RBCM rupture but allows PVM rupture and RBCM poration (Glushakova *et al*., 2010; Thomas *et al*., 2018). This showed that RBCM poration during egress was only subtly affected in the LCAT-null parasites (Figure 5F and G and Supplementary Movie S3). We further confirmed this subtle phenotype by quantifying porated parasitized RBCs in larger populations using flow cytometry (Figure 5H). We concluded that the egress defect could not be entirely due to a defect in RBCM poration.

### Phosphatidylserine and acylphosphatidylglycerol levels change during egress of LCAT-null parasites

Given the predicted role of LCAT as a catalytically active phospholipase, we reasoned that the LCAT-null phenotype likely resulted from a defect in parasite phospholipid modification prior to and/or during egress. To seek insights into this, we studied the phospholipid composition of LCAT-null mutants, comparing lipid profiles of RAP- and mock-treated LCAT:2loxPint schizonts just prior to egress.

Analysis by quantitative liquid chromatography-coupled mass spectrometry (LC-MS/MS) detected a total of 111 lipid species and found the lipid profiles of both schizont samples to be remarkably similar (Supplementary Figure S4A). This indicates that loss of LCAT does not have any detectable impact on the phospholipid composition during parasite development up to mature schizont stage.

Next, we profiled the phospholipid content of these schizonts before and immediately following egress to examine phospholipid-level changes occurring during egress (Figure 6A). For this, we first extracted lipids from synchronous, C2-arrested schizont populations of both RAP- and mock-treated parasites (time point before egress, BE). We then released the egress arrest by washing away the C2 and allowed the schizonts to undergo egress for 45 minutes in a small volume of culture media. The entire sample, confirmed by microscopy (Figure 6B) to comprise predominantly free merozoites, ruptured RBC and PV membranes, and the few residual schizonts that did not undergo egress, was subjected to lipid extraction (time point after egress, AE). Pairwise comparisons between the BE and AE samples within LCAT-null and wild type control parasites showed significant changes in abundance of several species belonging to three lipid classes-phosphatidylserine (PS), phosphatidylethanolamine (PE) and acylphosphatidylglycerol (acylPG) (Figure 6C and Supplementary Figure S4B). 9 out of 11 PS species detected were significantly enriched (1.5-2 fold) upon egress of the LCAT-null parasites but not during egress of wild type controls (Figure 6C and Supplementary Figure S5A). Similar enrichment was observed in some PE species in LCAT-null parasites while wild type egress inversely produced a significant decrease in several PE species. We also observed a striking decrease in all acylPG species (1.5 to 2-fold change) during egress of both LCAT-null and wild type schizonts which suggests that this decrease is normally associated with egress and is independent of LCAT activity (Figure 6C and Supplementary Figure S5B).

**Figure 6.**
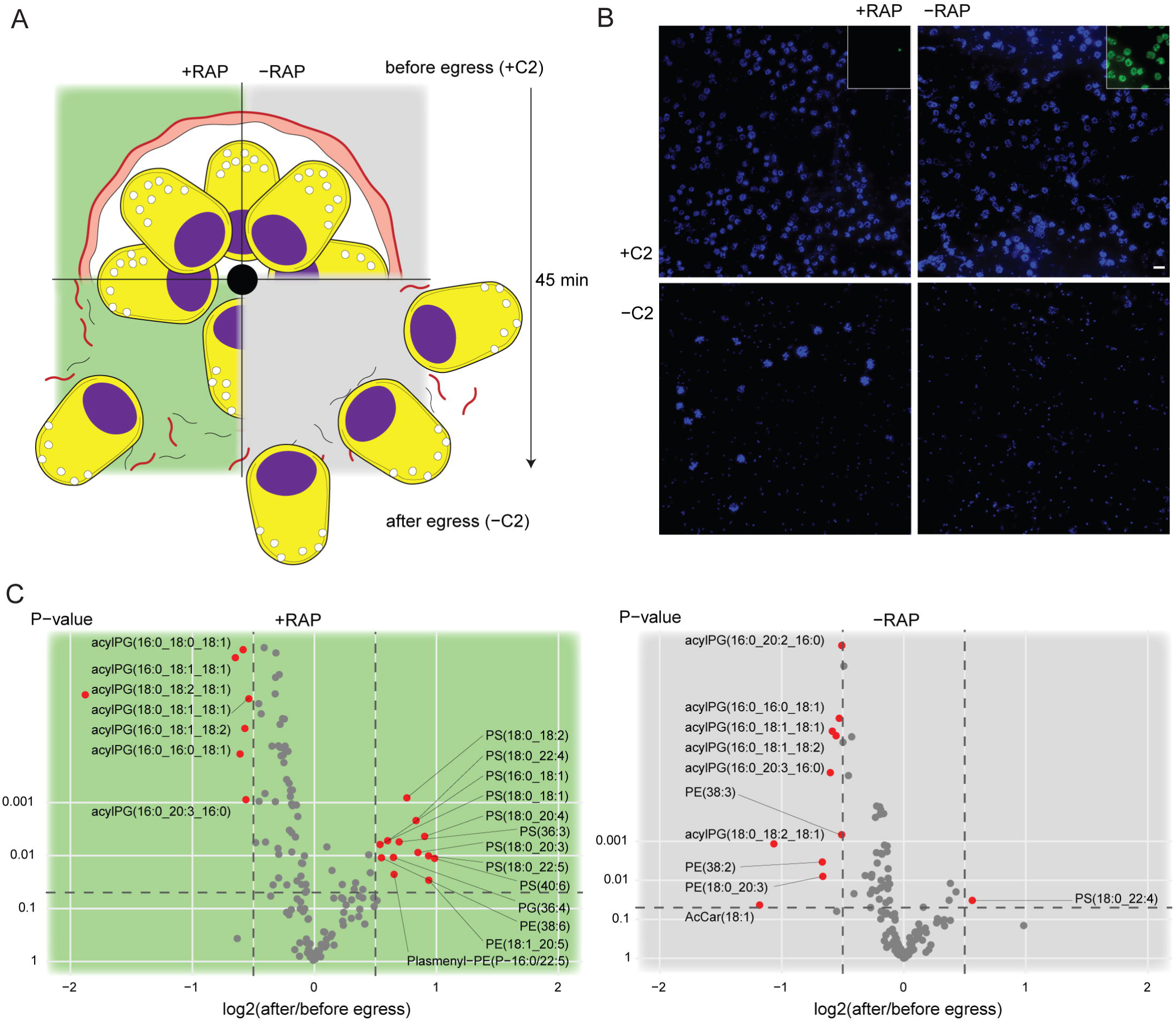
Lipidomic profiling of LCAT-null parasites reveals changes in phosphatidylserine and acylphosphatidylglycerol levels upon egress. **A)** Phospholipid content of RAP-treated (+RAP) and mock-treated (-RAP) LCAT:2loxPint parasites were assessed in egress-stalled schizonts (+C2) and after washing away C2, allowing egress to ensue for 45 min (-C2). -C2 suspensions contained free merozoites, remnants of ruptured PVM (black) and RBCM (red), a few un-egressed schizonts and in addition to this, merozoite clumps in the +RAP cultures due to inefficient egress. **B)** IFA confirming status of egress in +C2 and -C2 samples. The images show intact schizonts in +C2 samples and egressed merozoites (and merozoite clumps in the case of LCAT-null parasites) in -C2 samples. Inset, successful ablation of LCAT (green) expression in RAP-treated parasites. Nuclei were stained with DAPI (blue). Scale bar, 10 µm. **C)** Levels of acylphosphatidylglycerol (acylPG) decrease upon egress of both +RAP and -RAP parasites. An increase in several phosphatidylserine species is observed upon egress of LCAT-null parasites (comparisons were done across 6 independent egress experiments).

## Discussion

In the minutes leading to ABS parasite egress, the PV is the site of intense proteolytic and membranolytic activity. Secretory vesicles are discharged, the PV rounds up, and the PVM and RBCM are ruptured and porated respectively before final RBCM rupture. PVM rupture and RBCM poration are brought about by unidentified effector molecules that act within a very short time frame, and we speculate that both events may be mediated by the same effector molecule(s). To gain a better understanding of these events during egress, we here studied i) the role of two perforin-like proteins (PPLP1 and PPLP2); ii) a PV-resident phospholipase (LCAT); and iii) changes in protein and phospholipid content that occur during egress.

PPLP1 and PPLP2 have established roles in sporozoite egress from transient vacuoles (Risco-Castillo *et al*., 2015) and gametocyte egress from RBCs (Wirth *et al*., 2014) respectively. The potential role of these proteins in ABS egress, however, has been a matter of debate. Both proteins were shown to be detectable in *P. falciparum* asexual stages with suggested overlapping roles during blood stage egress (Garg *et al*., 2013) In contrast, in other studies, neither PLP was detected in ABS of rodent malaria parasites and individual null mutants exhibited normal blood stage growth (Deligianni *et al*., 2013; Ishino *et al*., 2005; Kaiser *et al*., 2004; Risco-Castillo *et al*., 2015; Wirth *et al*., 2014; Yang *et al*., 2017). However, individual gene knockouts do not rule out the possibility that the PLPs might play compensatory roles, a notion supported by the finding that small molecule PLP inhibitors have anti-parasite activity and block egress (Garg *et al*., 2020). We tested this hypothesis with a PPLP1/PPLP2 double knockout mutant. The PPLP1/PPLP2-null mutants exhibited normal growth rates, membrane poration and egress, confirming their combined dispensability in ABS growth. Our results contradict the findings of (Garg *et al*., 2020) and may suggest that the chemical inhibitors used in their study have off-target activity that is responsible for the observed egress block. Collectively, the evidence from all functional studies (including this study) of *Plasmodium* PLPs strongly suggests that these molecules have no important role in ABS parasite egress.

The malaria parasite secretes proteins into the PV through specialized apical organelles to perform functions that are vital for egress and invasion as well as modifying the nascent PV after invasion. It is becoming increasingly evident that subpopulations of these vesicles are discharged in a controlled way at specific times to perform specific functions (Absalon *et al*., 2018; Ebrahimzadeh et al., 2019). Our highly selective proteomics approach, comparing PV-extracts of SUB1-null and control parasites treated with an inhibitor of organelle discharge (C2), identified known microneme and exoneme proteins that are secreted into the PV within a short timeframe before egress. These include well-established micronemal proteins that are discharged onto the merozoite apical tip (MTRAP; (Baum et al., 2006)) or merozoite surface (GAMA, EBA-140 and EBA-181; (Arumugam et al., 2011; Gilberger et al., 2003; Hinds et al., 2009; Thompson et al., 2001)) during egress. Surprisingly no differences between the parasite extracts were observed in the levels of another micronemal *ebl* protein, EBA-175 (Reed et al., 2000), whereas the bona fide exoneme protease plasmepsin X (PMX) that proteolytically processes SUB1 was enriched in the C2-arrested extracts (Nasamu *et al*., 2017; Pino *et al*., 2017). In addition to this, we also found a putative nuclear export mediator factor (NEMF; PF3D7_1202600) enriched 2.5-fold that could potentially be studied further. This gene is continually expressed in ABS with peak transcription during schizont stages and is considered essential (Zhang et al., 2018). Our approach is by no means exhaustive and our workflow may have failed to capture low-abundance proteins. Our data, whilst not revealing new effector molecules, still essentially captured the PV proteome during egress and showed the presence of the known phospholipase LCAT in all our PV protein extracts, leading us to study this molecule further.

The role of LCAT has been previously examined in *Toxoplasma gondii* tachyzoites and *P. berghei* sporozoite and liver stages. LCAT orthologs in these organisms were found to be diversely localized depending on the parasite life stage. TgLCAT localizes to the plasma membrane in extracellular parasites and is secreted into the PV lumen late into the intracellular replication cycle via dense granule-like organelles (Pszenny et al., 2016; Schultz and Carruthers, 2018). Similarly, in *P. berghei,* PbLCAT is expressed on the surface of sporozoites (Bhanot et al., 2005) but after hepatocyte invasion mainly localizes to the PV and the PVM in addition to vesicular structures within the parasite cytoplasm (Burda *et al*., 2015). Loss of LCAT expression results in delayed egress of *T. gondii* tachyzoites from the host cell (Pszenny *et al*., 2016; Schultz and Carruthers, 2018), *P. berghei* sporozoites from oocysts, and *P. berghei* merozoites from liver-stage schizonts (Burda *et al*., 2015).

Here we have shown that LCAT localization and its facilitative role during egress observed elsewhere can now be extended to *P. falciparum* ABS schizonts too. We localised LCAT both to punctate signals within the cytoplasm (likely secretory organelles) as well as the PV lumen. Ablation of LCAT expression results in a reduced replication rate, caused by a defective egress phenotype in which RBCM poration is only mildly affected, RBCM rupture appears to progress normally, but merozoite dispersal is inefficient. We can infer from this that the PVM ruptures at least partially since the PV protein SERA6 is required to access the internal face of the RBCM for it to rupture. Partial or complete PVM rupture is also strongly supported by the fact that the egress defect can be overcome by mechanical shear stress, presumably helping to disperse the merozoites. In contrast, shear forces failed to rescue the egress defect in SUB1-null or SERA6-null schizonts where an intact RBCM and/or PVM was present (Thomas *et al*., 2018). Inefficient egress in LCAT-null liver stage schizonts was attributed to a defect in PVM disruption, albeit partial as a proportion of LCAT-null parasites were still able to disrupt the PVM (Burda *et al*., 2015). Our observations here indicate that inefficient egress in LCAT-null ABS schizonts cannot be attributed to a defect in PVM rupture.

Our results also contrast with the lack of phenotype observed in previous loss-of-function analyses in *P. berghei* and *P. falciparum* blood stages (Burda et al., 2021b; Burda *et al*., 2015). This is perhaps not surprising for *P. berghei* ABS as the *in vivo* environment (akin to mechanical shaking) likely ensures efficient dispersal of the LCAT-null merozoites. The ∼15% decrease in growth rate observed after conventional knockout of *P. falciparum* LCAT (as opposed to the ∼66% reduction we observed in our conditional knockout line) could be a result of parasite adaptation to in long-term culture. With the inducible DiCre system, we were able to clearly discern the LCAT-null phenotype by studying the mutants immediately after ablating LCAT expression, within the same erythrocytic cycle.

*P. berghei* LCAT has been shown to hydrolyse PC to produce lysoPC in vitro (at rates significantly lower than human LCAT) (Bhanot *et al*., 2005) which suggests that LCAT could lyse membranes either by direct hydrolysis of PC in the membrane or by producing lysoPC species which themselves are membranolytic (Bhanot *et al*., 2005; Pszenny *et al*., 2016). However, our extensive lipidomics analysis of LCAT-null schizonts showed no significant changes in PC or lysoPC levels either before or following egress. The nearly identical phospholipid profiles of LCAT-null and wildtype schizonts prior to egress either suggests that LCAT remains inactive before egress or that LCAT activity likely results in very small changes in phospholipid levels that cannot be discerned by our approach. By contrast, egress of both LCAT-null and wildtype schizonts was accompanied by distinct changes in phospholipid content. The accumulation of phosphatidylserine during LCAT-null schizont egress is intriguing. Phosphatidylserine is found significantly enriched in RBC-derived vesicles (RMVs) that are released from infected RBCs (Gulati et al., 2015). Release of RMVs peak during schizogony shortly before egress but do not occur during egress (Mantel et al., 2013). Therefore, it is plausible that inefficient egress of LCAT-null schizonts extend the period when RMVs are released thereby increasing PS content to a higher degree compared to wildtype schizonts. Unfortunately, our lipidomics efforts failed to provide any further insights into the exact role of LCAT in facilitating efficient egress.

The consistent depletion of acyl-phosphatidylglycerol (acyl-PG) over the course of ABS egress (irrespective of the presence or absence of LCAT) is intriguing. Acyl-PG (also known as semilysobisphosphatidic acid, its stereoisomeric counterpart) is an unusual glycerophospholipid with three acyl chains, one ester-bonded to the glycerol head group and the other two to the glycerol 3-phosphate backbone. The detected acyl-PG species possessed either palmitate (16:0) or stearate (18:0) at the head group while the other two acyl chains were combinations of saturated and unsaturated fatty acids. Acyl-PG was initially identified as a major phospholipid class in bacteria (Dalebroux et al., 2014; Yague et al., 1997) but has also been found enriched in Golgi membranes in rodents (Moreau et al., 2019) and is thought to play a role in membrane rupture and assembly during vaccinia virus assembly (Chlanda et al., 2009; Cluett and Machamer, 1996). Acyl-PG has been suggested to play a role in vesicle budding and fusion as its small polar head and three acyl chains gives the molecule a conical shape which can induce membrane curvature (Cluett et al., 1997; Zimmerberg and Kozlov, 2006). Phospholipases are known to interconvert cone-shaped PLs like acyl-PG and inverted cone shaped PLs (lysoPLs) to modulate membrane curvature during endocytosis (Brown et al., 2003). In *Plasmodium*, acyl-PG levels have been shown previously to peak in late schizonts (Gulati *et al*., 2015). Our results suggest that acyl-PG could be continuously delivered to the expanding PVM to maintain its curvature, then during egress a phospholipase degrades acyl-PG thereby causing the PVM to rupture. An alternative or additional hypothesis is that acyl-PG could be a major constituent of endocytic vesicles that deliver proteolytic enzymes to the PV prior to egress and their fusion to the PPM or PVM is facilitated by removal of acyl-PG by a phospholipase. We were unable to pursue these hypotheses further, in part because antibodies against acyl-PG suitable for localisation studies are unavailable and the molecular players in acyl-PG metabolism in general are also largely unknown. Recently, the phospholipase PfPATPL1 was found to play a role in gametocyte egress, with PfPATPL1-null parasites showing defects in rounding up and vesicular transport of proteins to the parasite periphery (Singh et al., 2019). While it is not known whether acyl-PG is deacylated by phospholipases, it is tempting to speculate that this phospholipid species plays an important role during egress.

How the PVM is ruptured during egress is a key question towards understanding the molecular mechanisms that bring about ABS egress. Whilst our diverse efforts here failed to answer this, we have eliminated PPLPs as prospective effectors, established a previously unknown role for the phospholipase LCAT and identified acyl-PG as an important phospholipid in asexual blood stage egress.

## Methods

### Plasmid construction

Modification plasmids to produce the five modified *P. falciparum* lines used in this study were constructed as follows.

The LCAT:GFP and LCAT:smMyc lines were made by tagging the endogenous *P. falciparum lcat* gene with GFP or smMyc using the selection-linked integration (SLI) method (Birnbaum *et al*., 2017). A GFP-tagging construct pSLI-PF3D7_0629300-GFP was generated by amplifying the C-terminal 900 bp of the *lcat* gene (PF3D7_0629300) using primers PF3D7_0629300-TAG-fw/ PF3D7_0629300-TAG-rev and cloning into pSLI-TGD (Birnbaum *et al*., 2017) using NotI/MluI.

Similarly, a smMyc-tagging construct pSLI-PF3D7_0629300-SM-Myc was generated by amplifying the smMyc sequence from pCAG-smFP Myc (Addgene plasmid #59757, gift from Loren Looger) (Viswanathan *et al*., 2015) using primer smMyc-fw/smMyc-rev and cloned into pSLI-PF3D7_0629300-GFP using MluI/SalI thereby replacing the GFP coding sequence with smMyc.

The conditional knockout lines were produced by modifying the endogenous target loci in the DiCre-expressing *P. falciparum* B11 line using Cas9-mediated genome editing (Ghorbal et al., 2014). A two-plasmid system was used where a targeting plasmid delivers Cas9 and guide RNA to target loci while a repair plasmid delivers the repair template for homology-directed repair of the Cas9-nicked locus.

The conditional double knockout PPLP1:loxNint/PPLP2:loxPint line was produced by sequentially floxing the endogenous *pplp1* (PF3D7_0408700) and *pplp2* (PF3D7_1216700) loci. Two RNA targeting sequences (TTTTAAAGCATTCTTAAATT for PPLP1 and TTTTTCTAGATATTCACCAA for PPLP2) were inserted into the pDC2 Cas9/gRNA/hDHFR (human dihydrofolate reductase)/yFCU (yeast cytosine deaminase/uridyl phosphoribosyl transferase)-containing plasmid as described previously (Knuepfer et al., 2017) to generate two different targeting plasmids (pCas9_pplp1_gRNA01 and pCas9_pplp2_gRNA01 respectively). For the repair plasmid pREP-PPLP1 for eight-exon *pplp1*, a recodonised segment of the coding region between second and sixth intron gene (628-2,577 bp; 99-587 aa) flanked by loxN-containing *sera2* introns (Jones et al., 2016) and ∼400-500 bp homology arms was synthesized commercially (GeneArt , Thermo Fisher Scientific) as a 2,361 bp long synthetic DNA fragment. Similarly, for pREP-PPLP2, the recodonised version of the *pplp2*’s MACPF domain (1,127-2,640 bp; 375-812 aa) flanked by loxPint modules (Jones *et al*., 2016) and ∼500 bp homology arms was synthesized commercially as a 2,497 bp long synthetic DNA fragment. Two different pairs of *lox* sites, the canonical loxP (core sequence-GCATACAT) and its variant loxN (core sequence - GTATACCT), were used for floxing to prevent cross-recombination events between both loci (Koussis et al., 2020). The repair plasmids were linearised overnight with PvuI and SacI prior to transfection.

The conditional frameshift-based knockout line LCAT:2loxPint, was produced by floxing a 200 bp region (973-1172bp) upstream of the 1,053 bp long catalytic domain that encodes an alpha/beta hydrolase fold with two catalytic GXSXG lipase motifs and a HX_4_D acyltransferase motif in the *lcat* gene (PF3D7_0629300). Two RNA targeting sequences (TAATAATAGAGATGAAATTT and ATAGAGATGAAATTTTGGTA) were inserted into the pDC2 Cas9/gRNA/hDHFR-containing plasmid to generate two different targeting plasmids (pCas9_lcat_gRNA01 and pCas9_lcat_gRNA02 respectively). For the repair plasmid pREP-LCAT, a 1,354 bp long synthetic DNA fragment containing a recodonised segment of the 200 bp flanked by 177 bp long loxPint modules and 400 bp homology arms was synthesized commercially. The repair plasmid was linearised with SphI overnight prior to transfection.

CloneAmp HiFi PCR Premix (TakaraBio) and Phusion High-Fidelity DNA polymerase (New England BioLabs) were used for PCR reactions for all plasmid constructions. All plasmid sequences were confirmed by Sanger sequencing. For sequences of oligonucleotides and other synthetic DNA used in this study, please refer to Supplementary File 1.

### Parasite culture maintenance, synchronisation and transfection

The DiCre-expressing *P. falciparum* B11 line (Perrin et al., 2018) was maintained at 37◦ C in human RBCs in RPMI 1640 containing Albumax II (Thermo Fisher Scientific) supplemented with 2 mM L-glutamine. Synchronisation of parasite cultures were done as described previously (Harris et al., 2005) by isolating mature schizonts by centrifugation over 70% (v/v) isotonic Percoll (GE Healthcare, Life Sciences) cushions, letting them rupture and invade fresh erythrocytes for 2 hours at 100rpm, followed by removal of residual schizonts by another Percoll separation and sorbitol treatment to finally obtain a highly synchronised preparation of newly invaded ring-stage parasites. To obtain the GDPD:HA:loxPint line, transfections were performed by introducing DNA into ∼10^8^ Percoll-enriched schizonts by electroporation using an Amaxa 4D Nucleofector X (Lonza), using program FP158 as previously described (Moon et al., 2013). For Cas9-based genetic modifications, 20 µg of targeting plasmid and 60 µg of linearised repair template were electroporated. Drug selection with 2.5 nM WR99210 was applied 24 h post-transfection for 4 days with successfully transfected parasites arising at 14-16 days. For sequential genetic modification, the PPLP1:loxPint line was treated with 1 μM 5-fluorocytosine (5-FC) provided as clinical grade Ancotil^®^ (MEDA) for one week before transfection with pCas9_pplp2_gRNA01 + pREP-PPLP2.

Clonal transgenic lines were obtained by serial limiting dilution in flat-bottomed 96-well plates (Thomas *et al*., 2016) followed by propagating wells that contain single plaques. Successful integration was confirmed by running diagnostic PCR either directly on culture using BloodDirect Phusion PCR premix or from extracted genomic DNA (DNAeasy Blood and Tissue kit, Qiagen) with CloneAmp HiFi PCR Premix (TakaraBio).

The *P. falciparum* 3D7 line was maintained at 37◦C in an atmosphere of 1% O_2_, 5% CO_2_, and 94% N_2_ and cultured using RPMI complete medium containing 0.5% Albumax according to standard procedures (Trager and Jensen, 1976). For generation of stable integrant cell lines LCAT:GFP and LCAT:smMyc, mature 3D7 schizonts were electroporated with 50 µg of plasmid DNA using a Lonza Nucleofector II device (Moon *et al*., 2013). and selected in medium supplemented with 3 nM WR99210 (Jacobus Pharmaceuticals). WR99210-resistant parasites were subsequently treated with 400 µg∕mL Neomycin/G418 (Sigma) to select for integrants carrying the desired genomic modification as described previously (Birnbaum *et al*., 2017). Successful integration was confirmed by diagnostic PCR using FIREpol DNA polymerase (Solis BioDyne).

To obtain LCAT-null parasites, DiCre-mediated excision of the target locus was induced by rapamycin treatment (100 nM RAP for 3 h or 10 nM overnight) of synchronous early ring-stage parasites (2–3 h post-invasion) as previously described (Collins et al., 2013a). Mock treated parasites were used as wild type controls.

### Western blot and immunofluorescence assays

To detect LCAT protein expression in LCAT:2loxPint line, rabbit polyclonal antibodies were raised against two peptide sequences within the N-terminal region of the *P. falciparum* LCAT protein; SIFLRNPYKITLGKSEK (32-48 aa) and FSEEEDSIVRRDTEKK (56-71 aa). For western blotting, proteins were extracted from C2-stalled mature schizonts directly into SDS buffer, resolved by SDS polyacrylamide gel electrophoresis (SDS-PAGE) and transferred to nitrocellulose membrane (Supported nitrocellulose membrane, Bio-Rad). Membranes were blocked with 5% bovine serum albumin (BSA) in PBS-T (0.05% Tween 20) and subsequently probed with the rabbit anti-LCAT sera (1:1,000 dilution), followed by horseradish peroxidase-conjugated goat anti-rabbit antibody (BioRad, 1:3,000). Immobilon Western Chemiluminescent HRP Substrate (Millipore) was used according to the manufacturer’s instructions, and blots were visualized and documented using a ChemiDoc Imager (Bio-Rad) with Image Lab software (Bio-Rad). Rabbit antibodies against SERA6 (Ruecker *et al*., 2012) was used at 1:1,000 as loading control.

For immunofluorescence assays of LCAT:2loxPint parasites, thin films of parasite cultures containing C2-arrested mature schizonts were air-dried, fixed in 4% (w/v) formaldehyde for 30 minutes (Agar Scientific Ltd.), permeabilized for 10 minutes in 0.1% (w/v) Triton ×100 and blocked overnight in 3% (w/v) bovine serum albumin in PBS. Slides were probed with rabbit anti-LCAT sera (1:1,000 dilution), human anti-MSP1 mAb X509 (1:1,000 dilution) (Blackman et al., 1991) and either mouse anti-AMA1-reduced (Collins et al., 2009) or mouse anti-SERA5 (Collins *et al*., 2017) (1:1,000 dilution). Primary antibodies were detected by probing with AlexaFluor 488-conjugated anti-rabbit, AlexaFluor 647-conjugated anti-human and AlexaFluor 594-conjugated anti-mouse (Invitrogen, 1:1,000) antibodies. Slides were then stained with 1ug/mL DAPI, mounted in Citifluor (Citifluor Ltd., Canterbury, U.K.).

For imaging LCAT:smMyc parasites, a circular space on a coverslip was surrounded with a Dako pen and coated with 10 μl 0.5 mg/ml Concanavalin A (ConA, in H_2_0, Sigma) in a humid chamber for 15 min at 37°C. 500 μl of Compound 2 (C2, 1 μM) arrested LCAT:smMyc schizont cultures were centrifuged and washed once in PBS to remove media components. ConA was washed away (3x H_2_0, 3x PBS) from coverslips and 50 μl of parasite cultures were applied to the coverslips and incubated at 37°C for 15 min in a humid chamber. Unbound cells were washed away with PBS and bound cells were fixed with 2% PFA / 0.0065% glutaraldehyde in PBS for 20 min at RT. Until this point, all solutions contained 1 μM C2 to prevent schizont egress. Cells were washed 3x with PBS and permeabilized with 0.2% Triton X-100 in PBS for 10 min at RT. Cells were then blocked for 10 minutes with 3% BSA/PBS followed by an incubation with 30 μl of primary antibody rabbit anti-Myc (1:1.000, Cell Signalling Technology, #2272) in 3% BSA/PBS in a humid chamber for 1 hour at RT. After washing cells 3x with PBS, cells were incubated in a humid chamber for 1 hour at RT with 30 μl of secondary antibody donkey anti-rabbit AlexaFluor488 (1:1.000, Invitrogen) in 3% BSA/PBS additionally containing 1 μg/ml DAPI for visualization of nuclei. After washing cells 3x with PBS, slides were mounted with Dako mounting solution, sealed and stored at 4°C protected from light until analysis.

### Fluorescence and time-lapse microscopy

For live cell microscopy of LCAT:GFP parasites, parasites were incubated with 1 µg∕mL DAPI in culture medium for 15 minutes at 37◦ C to stain nuclei before microscopic analysis. Parasites were imaged on a Leica D6B fluorescence microscope, equipped with a Leica DFC9000 GT camera and a Leica Plan Apochromat 100x/1.4 oil objective. Viewing chambers for live microscopy were constructed by adhering 22 × 64 mm borosilicate glass coverslips (VWR International) to microscope slides, as described previously (Collins *et al*., 2013a). Mature Percoll-enriched schizonts were incubated for ≥ 4 h at 37°C in Albumax-supplemented RPMI medium supplemented with 1 μM C2. Subsequently, ∼5 × 10^7^ schizonts were rapidly washed twice in 1 ml of gassed complete medium pre-warmed to 37°C and lacking C2, pelleting at 1,800 × *g* for 1 min. The cells were suspended in 60 μl of the same medium and introduced into a pre-warmed viewing chamber which was then immediately placed on a temperature-controlled microscope stage held at 37°C on a Nikon Eclipse Ni-E wide-field microscope fitted with a Nikon N Plan Apo λ 100×/1.45NA oil immersion objective and a Hamamatsu C11440 digital camera and documented via the NIS Elements software (Nikon). Images were acquired at 5- to 10-s intervals over a total of 30-40 min then processed and exported as TIFFs using Fiji (Schneider et al., 2012). For simultaneous capture of egress of RAP- and mock-treated schizonts in the same chamber, one of the lines were stained with 1ug/uL Hoechst in C2-supplemented media at 37°C for 5 minutes and washed twice with 1 mL C2-supplemented media before adding to an equal amount of unstained line and proceeding as before. RAP- and mock-treated lines were stained alternatively for each video. For visualising RBCM poration, schizonts were washed and then suspended in 60 μL media with E64 (50 μM final concentration) and AlexaFluor 488 phalloidin (Invitrogen; diluted 1:50 from 200 unit ml^-1^ stock in PBS) and proceeded as before for imaging. For flow cytometry-based quantification of porated RBCs, egress-arrested schizonts were stained with Hoechst in C2-supplemented media at 37°C for 5 minutes and washed once with 1 mL C2-supplemented media. Hoechst-stained schizonts were washed and then resuspended in either 200uL media with E64 and AlexaFluor 488 phalloidin and allowed to egress for half hour at 37°C in tubes. As a negative control, another aliquot of the schizonts were incubated in the presence of C2 instead of E64. Samples before and after incubation (t = 0 and t = 30min) were analysed by flow cytometry on a BD FACSVerse using BD FACSuite software. For every sample, 10,000 - 30,000 events were recorded and filtered with appropriate forward and side scatter parameters. Hoechst-positive (infected RBCs) were gated using a 448/45 detector configuration and of this, AlexaFluor 488 phalloidin-positive RBCs were counted using a 527/32 detector configuration. All data were analysed using FlowJo software.

### Growth and replication assays

Growth assays were performed to assess parasite growth across 3-4 erythrocytic replication cycles. Synchronous cultures of ring-stage parasites at 0.1% parasitaemia and 2% haematocrit were maintained in triplicates in 12 well plates. 50 µL from each well was sampled at 0, 2, 4 and 6 days post-RAP treatment, fixed with 50 µL of 0.2% glutaraldehyde in PBS and stored at 4◦ C for flow cytometry quantification. Fixed parasites were stained with SYBR Green (Thermo Fisher Scientific, 1:10,000 dilution) for 20 min at 37◦ C and analysed by flow cytometry on a BD FACSVerse using BD FACSuite software. For every sample, parasitaemia was estimated by recording 10,000 events and filtering with appropriate forward and side scatter parameters and gating for SYBR Green stain-positive (infected RBCs) and negative RBCs using a 527/32 detector configuration. All data were analysed using FlowJo software. Growth stage progression was monitored by microscopic examination at selected timepoints using Giemsa-stained thin blood films.

Plaque growth assays were performed by dispensing around 20 parasites per well in flat-bottomed microplates at a haematocrit of 0.75%, as described (Thomas *et al*., 2016). Plates were imaged using a high resolution flat-bed scanner 14–16 days after setting up the assays. Plaques were counted by visual examination of the images and plaque size quantified using the Lasso tool in Adobe Photoshop CS6.

To assess invasion rates, highly synchronous mature schizonts were added to fresh erythrocytes (2% haematocrit) and let to invade for four hours at both static and mechanical shaking (100 rpm) conditions (four replicates in each condition). Cultures were sampled before and after the 4 h invasion and fixed as before for quantification.

### Proteomic analysis

To assess the changes in PV proteome upon micronemal/exonemal release, synchronous SUB1HA3:loxP parasites (Thomas *et al*., 2018) at ∼32 hpi were treated with 1 μM C2 overnight. To trigger micronemal/exonemal release, schizonts were washed with RPMI w/o Albumax and incubated in RPMI w/o Albumax for 20 min at 37° C. C2-treated and C2-washed schizonts were lysed in ice cold 0.15% saponin (with the addition of C2 for the +C2 samples). The saponin fractions were filtered using Ultrafree-MC 0.22 μm GV Durapore (Milipore) filters spun at 13,000 rpm for 1 min. Multiple aliquots of the saponin fractions were snap frozen on dry ice/ethanol. Proteins were denatured by adding 20 μl of 4x Laemmli buffer with 10 mM DTT freshly added and heating at 95° C for 5 min. Denatured proteins were run on a BioRad TGX 4-15 % Tris-Glycine gel and then proceeded with in-gel digestion.

Reduced and alkylated proteins were in-gel digested with 100ng trypsin (modified sequencing grade, Promega) overnight at 37° C. Supernatants were dried in a vacuum centrifuge and resuspended in 0.1% TriFluoroAcetic acid (TFA).

On an Ultimate 3000 nanoRSLC HPLC (Thermo Scientific) 1-10ul of acidified protein digest was loaded onto a 20mm × 75um Pepmap C18 trap column (Thermo Scientific) prior to elution via a 50cm × 75um EasySpray C18 column into a Lumos Tribrid Orbitrap mass spectrometer (Thermo Scientific). A 90’ binary gradient of 6%-40%B over 63’ was used prior to washing and re-equilibration (A= 2%ACN, 0.1% formic acid; B= 80%ACN, 0.1% formic acid).

The Orbitrap was operated in ‘TopS’ Data Dependent Acquisition mode with precursor ion spectra acquired at 120k resolution in the Orbitrap detector and MS/MS spectra at 32% HCD collision energy in in the ion trap. Automatic Gain Control was set to Auto for MS1 and MS2. Maximum injection times were set to ‘Standard’ (MS1) and ‘Dynamic’ (MS2). Dynamic exclusion was set to 20s.

Raw files were processed using Maxquant (maxquant.org) and Perseus (maxquant.net/perseus) with recent downloads of the Plasmodium falciparum 3D7 (www.plasmodb.org) and the Uniprot Homo sapiens reference proteome, together with the Maxquant common contaminants databases. A decoy database of reversed sequences was used to filter false positives at protein and peptide FDR of 1%. T-tests were performed with a permutation-based FDR of 5% to cater for multiple hypothesis testing.

### Lipidomic analysis

To assess the changes in phospholipid content due to absence of LCAT, total phospholipids from LCAT-null and wildtype schizonts were extracted and lipid species were determined and quantified by LC-MS/MS. Schizonts were isolated using Percoll cushions from RAP- and mock-treated LCAT:2loxPint parasitized cultures (100ml, 0.5% haematocrit, 35-40% parasitaemia) grown for 45 hours post treatment and allowed to mature for 4 hours at 37◦ C in the presence of egress-blocking C2 (1 µM) in order to achieve a high level of homogeneity in the samples. Egress-blocked schizonts were washed twice with RPMI media w/o Albumax II (with C2 at 1 µM) and subject to lipid extraction. Lipid extraction for each sample was performed by adding 400 µL of 1 × 10^8^ parasites, either as intact schizonts or egressed suspensions, to each of three tubes (technical replicates) that contained 600 µL methanol and 200 µL chloroform. Experiments were carried out in triplicate. To assess the changes in phospholipid content upon egress, egress-blocked LCAT-null (+RAP) and wildtype (-RAP) schizonts (in RPMI media without Albumax II with C2 at 1 µM) were divided into 12 aliquots and kept at 37◦ C. Of these, 6 aliquots were washed with prewarmed RPMI media w/o Albumax and incubated in 200uL prewarmed RPMI media w/o Albumax at 37◦ C for 45 min for them to egress. Total phospholipids were extracted from the six egress-blocked samples (+C2) and from the six independently egressed samples (-C2) by adding 200 µL of approximately 1 × 10^10^ parasites, either as intact schizonts or egressed suspensions, to 600 µL methanol and 200 µL chloroform. Samples were sonicated for 8 minutes at 4◦ C and incubated at 4◦ C for 1 hour. 400 µL of ice-cold water was added (thus obtaining the 3:3:1 water:methanol:chloroform ratio) to the samples, mixed well and centrifuged at max speed for 5 min at 4◦ C for biphasic partitioning. The lower apolar phase was added to fresh tubes. The upper aqueous layer was removed and lipids were extracted once more by adding 200 µL of chloroform, vortexing and centrifuging as before. The apolar phases from both extractions were pooled (400 µL) and dried under nitrogen stream and resuspended in butanol/methanol (1:1,v/v) containing 5 µM ammonium formate.

The LC-MS method was adapted from (Greenwood et al., 2019). Cellular lipids were separated by injecting 10 µL aliquots onto a column: 2.1 × 100 mm, 1.8 µm C18 Zorbax Elipse plus column (Agilent) using an Dionex UltiMate 3000 LC system (Thermo Scientific). A 20 min elution gradient of 45% to 100% Solvent B was used, followed by a 5 min wash of 100% Solvent B and 3 min re-equilibration, where Solvent B was water:acetonitrile:isopropanol, 5:20:75 (v/v/v) with 10 mM ammonium formate (Optima HPLC grade, Fisher Chemical) and Solvent A was 10 mM ammonium formate in water (Optima HPLC grade, Fisher Chemical). Other parameters were as follows: flow rate 600 µL /min; column temperature 60◦ C; autosampler temperature 10◦ C. MS was performed with positive/negative polarity switching using an Q Exactive Orbitrap (Thermo Scientific) with a HESI II probe. MS parameters were as follows: spray voltage 3.5 kV and 2.5 kV for positive and negative modes, respectively; probe temperature 275◦ C; sheath and auxiliary gases were 55 and 15 arbitrary units, respectively; full scan range: 150 to 2000 m/z with settings of auto gain control (AGC) target and resolution as Balanced and High (3 × 10^6^ and 70,000), respectively. Data was recorded using Xcalibur 3.0.63 software (Thermo Scientific). Mass calibration was performed for both ESI polarities before analysis using the standard Thermo Scientific Calmix solution. To enhance calibration stability, lock-mass correction was also applied to each analytical run using ubiquitous low-mass contaminants. To confirm the identification of significant features, pooled quality control samples were ran in data-dependent top-N (ddMS2-topN) mode, acquisition parameters as follows: resolution of 17,500, auto gain control target under 2 × 10^5^, isolation window of m/z 0.4 and stepped collision energy 10, 20 and 30 in HCD (high-energy collisional dissociation) mode. Data analysis was performed using Free Style 1.5 (ThermoScientific), Progenesis (Nonlinear Dynamics) and LipidMatch (Koelmel et al., 2017).

### Statistical analysis

All statistical analysis and data visualization was performed in R v4.0.2 (R Core Team (2021)). Unless stated otherwise, Student’s t-test were used to compare group means and where necessary Bonferroni adjustment for multiple comparisons was applied to the p-value of statistical significance. All statistical analysis is available as R code in https://github.com/a2g1n/LCATxcute.

## Author contributions

Conceptualization: AR, PCB, KK, TWG and MJB. Investigation and Methodology: AR, PCB, KK, JAT, EP, EC, SAH, JIM, APS, TWG and MJB. Data curation and Formal analysis: AR, PCB, EC and SAH. Visualization and Writing – original draft: AR. Writing-review & editing: AR, PCB, TWG and MJB. Funding acquisition and Supervision: TWG and MJB.

## Acknowledgement

AR was funded by a Marie Skłodowska Curie Individual Fellowship (Project number 751865). The work was also supported by funding to MJB from the Wellcome Trust (20318/A/20/Z) and the Francis Crick Institute (https://www.crick.ac.uk/) which receives its core funding from Cancer Research UK (CC2129), the UK Medical Research Council (CC2129), and the Wellcome Trust (CC2129). For the purpose of Open Access, the author has applied a CC BY public copyright licence to any Author Accepted Manuscript version arising from this submission. The work was further supported by Wellcome ISSF2 funding to the London School of Hygiene and Tropical Medicine. PCB acknowledges funding by the German Research Foundation (DFG project number 414222880). Images were acquired on microscopes of the CSSB imaging facility.

---

**Supplementary Figure 1. A)** Diagnostic PCR showing correct integration of the modification plasmids into the PPLP1 and PPLP2 loci in PPLP1:loxNint/PPLP2:loxPint parasites. Primers used are denoted in Figure 1A. **B)** Replication of PLP1:loxPint clonal line prior to second modification. The modified parasites show a normal replication rate across two cycles (error bars, ± SD, triplicate RAP treatments with different blood sources).

**Supplementary Figure S2. A)** Strategy for SLI-based endogenous tagging of the *lcat* gene with GFP. Primers used for integration PCR are indicated with half arrows. T2A, skip peptide; Neo-R, neomycin-resistance gene; hDHFR, human dihydrofolate reductase; lollipop, stop codons; arrows, promoters. **B)** Diagnostic PCR showing correct integration of the modification plasmid into the LCAT locus in the LCAT:GFP parasites. KI, knock in cell line; WT, wild type parental line. **C)** Strategy for SLI-based endogenous tagging of *lcat* gene with smMyc. Primers used for integration PCR are indicated with half arrows. **D)** Diagnostic PCR showing correct integration of the modification plasmid into the LCAT locus in the LCAT:smMyc line. **E)** IFA of LCAT:smMyc mature schizonts using anti-myc (green) antibodies. DAPI-stained nuclei are shown in blue. DIC, differential interference contrast. Scale bar, 5 µm.

**Supplementary Figure S3. A)** Diagnostic PCR showing correct integration of the modification plasmid into the LCAT locus in LCAT:2loxPint line. Primers used are indicated in Figure 4A.

**Supplementary Figure S4. A)** Lipidomic analysis of LCAT:2loxPint egress-stalled schizonts following mock-or RAP-treatment at ring stages. The bubble plot shows the fold change in levels of various lipid species in LCAT-null schizonts compared to controls (3 independent biological replicates). No significant change in phospholipid levels were detected between the samples. **B)** Bubble plot shows the fold change in levels of various lipid species before and after egress of RAP-treated (+RAP) and mock-treated (-RAP) LCAT:2loxPint parasites (6 independent biological replicates).

**Supplementary Figure S5.** Relative peak intensities and log_2_ fold change (depicted as dot plots) of the significantly altered **A)** phosphatidylserine and **B)** acylphosphatidylglycerol species upon egress of mock- or RAP-treated LCAT:2loxPint schizonts. B) comparison between choline-starved GDPD:loxPint:HA (B4) and PfGDPD-null (clone G1) parasites.

**Supplementary Table S1. Sequences of oligonucleotides and other synthetic DNA used in this study.**

**Supplementary Table S2. Label-free quantitation of proteins detected in SUB1-null schizonts in the presence or absence of C2.**

**Supplementary Table S3. Raw peak intensities of various lipid species measured in LCAT:2loxPint egress-stalled schizonts following mock-or RAP-treatment at ring stages.**

**Supplementary Table S4. Raw peak intensities of various lipid species measured before and after egress in RAP-treated (+RAP) and mock-treated (-RAP) LCAT:2loxPint parasites.**

**Supplementary movie S1. Composite time-lapse video showing RAP- and mock-treated (blue; stained with DAPI) PPLP1:loxNint/PPLP2:loxPint parasites undergoing normal RBCM poration and egress as visualized with fluorescent phalloidin (green) that gains access and binds to the RBC cytoskeleton upon RBCM poration.**

**Supplementary movie S2. Composite time-lapse video showing different fates of RAP- (blue; stained with Hoechst) and mock-treated LCAT:2loxPint parasites.** More number of abnormal egress events (marked with red circles; normal egress events in green) were observed in RAP-treated LCAT:2loxPint parasites compared to mock-treated parasites.

**Supplementary movie S3. Composite time-lapse video showing normal RBCM poration in both RAP-(blue; stained with Hoechst) and mock-treated LCAT:2loxPint parasites as visualized with fluorescent phalloidin (green).**

## References

Abkarian, M., Massiera, G., Berry, L., Roques, M., and Braun-Breton, C. (2011). A novel mechanism for egress of malarial parasites from red blood cells. Blood 117, 4118–4124. 10.1182/blood-2010-08-299883.

Absalon, S., Blomqvist, K., Rudlaff, R.M., DeLano, T.J., Pollastri, M.P., and Dvorin, J.D. (2018). Calcium-Dependent Protein Kinase 5 Is Required for Release of Egress-Specific Organelles in Plasmodium falciparum. mBio 9. 10.1128/mBio.00130-18.

Arumugam, T.U., Takeo, S., Yamasaki, T., Thonkukiatkul, A., Miura, K., Otsuki, H., Zhou, H., Long, C.A., Sattabongkot, J., Thompson, J., et al. (2011). Discovery of GAMA, a Plasmodium falciparum merozoite micronemal protein, as a novel blood-stage vaccine candidate antigen. Infect Immun 79, 4523–4532. 10.1128/IAI.05412-11.

Baum, J., Richard, D., Healer, J., Rug, M., Krnajski, Z., Gilberger, T.W., Green, J.L., Holder, A.A., and Cowman, A.F. (2006). A conserved molecular motor drives cell invasion and gliding motility across malaria life cycle stages and other apicomplexan parasites. J Biol Chem 281, 5197–5208. 10.1074/jbc.M509807200.

Bhanot, P., Schauer, K., Coppens, I., and Nussenzweig, V. (2005). A surface phospholipase is involved in the migration of plasmodium sporozoites through cells. J Biol Chem 280, 6752–6760. 10.1074/jbc.M411465200.

Birnbaum, J., Flemming, S., Reichard, N., Soares, A.B., Mesen-Ramirez, P., Jonscher, E., Bergmann, B., and Spielmann, T. (2017). A genetic system to study Plasmodium falciparum protein function. Nat Methods 14, 450–456. 10.1038/nmeth.4223.

Blackman, M.J., Whittle, H., and Holder, A.A. (1991). Processing of the Plasmodium falciparum major merozoite surface protein-1: identification of a 33-kilodalton secondary processing product which is shed prior to erythrocyte invasion. Mol Biochem Parasitol 49, 35–44. 10.1016/0166-6851(91)90128-s.

Brown, W.J., Chambers, K., and Doody, A. (2003). Phospholipase A2 (PLA2) enzymes in membrane trafficking: mediators of membrane shape and function. Traffic 4, 214–221. 10.1034/j.1600-0854.2003.00078.x.

Burda, P.-C., Ramaprasad, A., Pietsch, E., Bielfeld, S., Söhnchen, C., Wilcke, L., Strauss, J., Schwudke, D., Sait, A., Collinson, L., et al. (2021a). Global analysis of putative phospholipases in the malaria parasite Plasmodium falciparum reveals critical factors for parasite proliferation. bioRxiv.

Burda, P.-C., Ramaprasad, A., Pietsch, E., Bielfeld, S., Söhnchen, C., Wilcke, L., Strauss, J., Schwudke, D., Sait, A., Collinson, L.M., et al. (2021b). Global analysis of putative phospholipases in the malaria parasite *Plasmodium falciparum* reveals critical factors for parasite proliferation. bioRxiv, 2021.2006.2028.450158. 10.1101/2021.06.28.450158.

Burda, P.C., Roelli, M.A., Schaffner, M., Khan, S.M., Janse, C.J., and Heussler, V.T. (2015). A Plasmodium phospholipase is involved in disruption of the liver stage parasitophorous vacuole membrane. PLoS Pathog 11, e1004760. 10.1371/journal.ppat.1004760.

Chlanda, P., Carbajal, M.A., Cyrklaff, M., Griffiths, G., and Krijnse-Locker, J. (2009). Membrane rupture generates single open membrane sheets during vaccinia virus assembly. Cell Host Microbe 6, 81–90. 10.1016/j.chom.2009.05.021.

Cluett, E.B., Kuismanen, E., and Machamer, C.E. (1997). Heterogeneous distribution of the unusual phospholipid semilysobisphosphatidic acid through the Golgi complex. Mol Biol Cell 8, 2233–2240. 10.1091/mbc.8.11.2233.

Cluett, E.B., and Machamer, C.E. (1996). The envelope of vaccinia virus reveals an unusual phospholipid in Golgi complex membranes. J Cell Sci 109 ( Pt 8), 2121–2131.

Collins, C.R., Das, S., Wong, E.H., Andenmatten, N., Stallmach, R., Hackett, F., Herman, J.P., Muller, S., Meissner, M., and Blackman, M.J. (2013a). Robust inducible Cre recombinase activity in the human malaria parasite Plasmodium falciparum enables efficient gene deletion within a single asexual erythrocytic growth cycle. Mol Microbiol 88, 687–701. 10.1111/mmi.12206.

Collins, C.R., Hackett, F., Atid, J., Tan, M.S.Y., and Blackman, M.J. (2017). The Plasmodium falciparum pseudoprotease SERA5 regulates the kinetics and efficiency of malaria parasite egress from host erythrocytes. PLoS Pathog 13, e1006453. 10.1371/journal.ppat.1006453.

Collins, C.R., Hackett, F., Strath, M., Penzo, M., Withers-Martinez, C., Baker, D.A., and Blackman, M.J. (2013b). Malaria parasite cGMP-dependent protein kinase regulates blood stage merozoite secretory organelle discharge and egress. PLoS Pathog 9, e1003344. 10.1371/journal.ppat.1003344.

Collins, C.R., Withers-Martinez, C., Hackett, F., and Blackman, M.J. (2009). An inhibitory antibody blocks interactions between components of the malarial invasion machinery. PLoS Pathog 5, e1000273. 10.1371/journal.ppat.1000273.

Dal Peraro, M., and van der Goot, F.G. (2016). Pore-forming toxins: ancient, but never really out of fashion. Nat Rev Microbiol 14, 77–92. 10.1038/nrmicro.2015.3.

Dalebroux, Z.D., Matamouros, S., Whittington, D., Bishop, R.E., and Miller, S.I. (2014). PhoPQ regulates acidic glycerophospholipid content of the Salmonella Typhimurium outer membrane. Proc Natl Acad Sci U S A 111, 1963–1968. 10.1073/pnas.1316901111.

Das, S., Hertrich, N., Perrin, A.J., Withers-Martinez, C., Collins, C.R., Jones, M.L., Watermeyer, J.M., Fobes, E.T., Martin, S.R., Saibil, H.R., et al. (2015). Processing of Plasmodium falciparum Merozoite Surface Protein MSP1 Activates a Spectrin-Binding Function Enabling Parasite Egress from RBCs. Cell Host Microbe 18, 433–444. 10.1016/j.chom.2015.09.007.

Deligianni, E., Morgan, R.N., Bertuccini, L., Wirth, C.C., Silmon de Monerri, N.C., Spanos, L., Blackman, M.J., Louis, C., Pradel, G., and Siden-Kiamos, I. (2013). A perforin-like protein mediates disruption of the erythrocyte membrane during egress of Plasmodium berghei male gametocytes. Cell Microbiol 15, 1438–1455. 10.1111/cmi.12131.

Ebrahimzadeh, Z., Mukherjee, A., Crochetiere, M.E., Sergerie, A., Amiar, S., Thompson, L.A., Gagnon, D., Gaumond, D., Stahelin, R.V., Dacks, J.B., and Richard, D. (2019). A pan-apicomplexan phosphoinositide-binding protein acts in malarial microneme exocytosis. EMBO Rep 20. 10.15252/embr.201847102.

Garg, S., Agarwal, S., Kumar, S., Yazdani, S.S., Chitnis, C.E., and Singh, S. (2013). Calcium-dependent permeabilization of erythrocytes by a perforin-like protein during egress of malaria parasites. Nat Commun 4, 1736. 10.1038/ncomms2725.

Garg, S., Shivappagowdar, A., Hada, R.S., Ayana, R., Bathula, C., Sen, S., Kalia, I., Pati, S., Singh, A.P., and Singh, S. (2020). Plasmodium Perforin-Like Protein Pores on the Host Cell Membrane Contribute in Its Multistage Growth and Erythrocyte Senescence. Front Cell Infect Microbiol 10, 121. 10.3389/fcimb.2020.00121.

Ghorbal, M., Gorman, M., Macpherson, C.R., Martins, R.M., Scherf, A., and Lopez-Rubio, J.J. (2014). Genome editing in the human malaria parasite Plasmodium falciparum using the CRISPR-Cas9 system. Nat Biotechnol 32, 819–821. 10.1038/nbt.2925.

Gilberger, T.W., Thompson, J.K., Reed, M.B., Good, R.T., and Cowman, A.F. (2003). The cytoplasmic domain of the Plasmodium falciparum ligand EBA-175 is essential for invasion but not protein trafficking. J Cell Biol 162, 317–327. 10.1083/jcb.200301046.

Glushakova, S., Beck, J.R., Garten, M., Busse, B.L., Nasamu, A.S., Tenkova-Heuser, T., Heuser, J., Goldberg, D.E., and Zimmerberg, J. (2018). Rounding precedes rupture and breakdown of vacuolar membranes minutes before malaria parasite egress from erythrocytes. Cell Microbiol 20, e12868. 10.1111/cmi.12868.

Glushakova, S., Humphrey, G., Leikina, E., Balaban, A., Miller, J., and Zimmerberg, J. (2010). New stages in the program of malaria parasite egress imaged in normal and sickle erythrocytes. Curr Biol 20, 1117–1121. 10.1016/j.cub.2010.04.051.

Glushakova, S., Yin, D., Li, T., and Zimmerberg, J. (2005). Membrane transformation during malaria parasite release from human red blood cells. Curr Biol 15, 1645–1650. 10.1016/j.cub.2005.07.067.

Greenwood, D.J., Dos Santos, M.S., Huang, S., Russell, M.R.G., Collinson, L.M., MacRae, J.I., West, A., Jiang, H., and Gutierrez, M.G. (2019). Subcellular antibiotic visualization reveals a dynamic drug reservoir in infected macrophages. Science 364, 1279–1282. 10.1126/science.aat9689.

Guerra, A.J., and Carruthers, V.B. (2017). Structural Features of Apicomplexan Pore-Forming Proteins and Their Roles in Parasite Cell Traversal and Egress. Toxins (Basel) 9. 10.3390/toxins9090265.

Gulati, S., Ekland, E.H., Ruggles, K.V., Chan, R.B., Jayabalasingham, B., Zhou, B., Mantel, P.Y., Lee, M.C., Spottiswoode, N., Coburn-Flynn, O., et al. (2015). Profiling the Essential Nature of Lipid Metabolism in Asexual Blood and Gametocyte Stages of Plasmodium falciparum. Cell Host Microbe 18, 371–381. 10.1016/j.chom.2015.08.003.

Hale, V.L., Watermeyer, J.M., Hackett, F., Vizcay-Barrena, G., van Ooij, C., Thomas, J.A., Spink, M.C., Harkiolaki, M., Duke, E., Fleck, R.A., et al. (2017). Parasitophorous vacuole poration precedes its rupture and rapid host erythrocyte cytoskeleton collapse in Plasmodium falciparum egress. Proc Natl Acad Sci U S A 114, 3439–3444. 10.1073/pnas.1619441114.

Harris, P.K., Yeoh, S., Dluzewski, A.R., O’Donnell, R.A., Withers-Martinez, C., Hackett, F., Bannister, L.H., Mitchell, G.H., and Blackman, M.J. (2005). Molecular identification of a malaria merozoite surface sheddase. PLoS Pathog 1, 241–251. 10.1371/journal.ppat.0010029.

Hinds, L., Green, J.L., Knuepfer, E., Grainger, M., and Holder, A.A. (2009). Novel putative glycosylphosphatidylinositol-anchored micronemal antigen of Plasmodium falciparum that binds to erythrocytes. Eukaryot Cell 8, 1869–1879. 10.1128/EC.00218-09.

Hybiske, K., and Stephens, R.S. (2008). Exit strategies of intracellular pathogens. Nat Rev Microbiol 6, 99–110. 10.1038/nrmicro1821.

Ishino, T., Chinzei, Y., and Yuda, M. (2005). A Plasmodium sporozoite protein with a membrane attack complex domain is required for breaching the liver sinusoidal cell layer prior to hepatocyte infection. Cell Microbiol 7, 199–208. 10.1111/j.1462-5822.2004.00447.x.

Jones, M.L., Das, S., Belda, H., Collins, C.R., Blackman, M.J., and Treeck, M. (2016). A versatile strategy for rapid conditional genome engineering using loxP sites in a small synthetic intron in Plasmodium falciparum. Sci Rep 6, 21800. 10.1038/srep21800.

Kaiser, K., Camargo, N., Coppens, I., Morrisey, J.M., Vaidya, A.B., and Kappe, S.H. (2004). A member of a conserved Plasmodium protein family with membrane-attack complex/perforin (MACPF)-like domains localizes to the micronemes of sporozoites. Mol Biochem Parasitol 133, 15–26. 10.1016/j.molbiopara.2003.08.009.

Khosh-Naucke, M., Becker, J., Mesen-Ramirez, P., Kiani, P., Birnbaum, J., Frohlke, U., Jonscher, E., Schluter, H., and Spielmann, T. (2018). Identification of novel parasitophorous vacuole proteins in P. falciparum parasites using BioID. Int J Med Microbiol 308, 13–24. 10.1016/j.ijmm.2017.07.007.

Knuepfer, E., Napiorkowska, M., van Ooij, C., and Holder, A.A. (2017). Generating conditional gene knockouts in Plasmodium - a toolkit to produce stable DiCre recombinase-expressing parasite lines using CRISPR/Cas9. Sci Rep 7, 3881. 10.1038/s41598-017-03984-3.

Koelmel, J.P., Kroeger, N.M., Ulmer, C.Z., Bowden, J.A., Patterson, R.E., Cochran, J.A., Beecher, C.W.W., Garrett, T.J., and Yost, R.A. (2017). LipidMatch: an automated workflow for rule-based lipid identification using untargeted high-resolution tandem mass spectrometry data. BMC Bioinformatics 18, 331. 10.1186/s12859-017-1744-3.

Koussis, K., Withers-Martinez, C., Baker, D.A., and Blackman, M.J. (2020). Simultaneous multiple allelic replacement in the malaria parasite enables dissection of PKG function. Life Sci Alliance 3. 10.26508/lsa.201900626.

Koussis, K., Withers-Martinez, C., Yeoh, S., Child, M., Hackett, F., Knuepfer, E., Juliano, L., Woehlbier, U., Bujard, H., and Blackman, M.J. (2009). A multifunctional serine protease primes the malaria parasite for red blood cell invasion. EMBO J 28, 725–735. 10.1038/emboj.2009.22.

Lopez-Barragan, M.J., Lemieux, J., Quinones, M., Williamson, K.C., Molina-Cruz, A., Cui, K., Barillas-Mury, C., Zhao, K., and Su, X.Z. (2011). Directional gene expression and antisense transcripts in sexual and asexual stages of Plasmodium falciparum. BMC Genomics 12, 587. 10.1186/1471-2164-12-587.

Mantel, P.Y., Hoang, A.N., Goldowitz, I., Potashnikova, D., Hamza, B., Vorobjev, I., Ghiran, I., Toner, M., Irimia, D., Ivanov, A.R., et al. (2013). Malaria-infected erythrocyte-derived microvesicles mediate cellular communication within the parasite population and with the host immune system. Cell Host Microbe 13, 521–534. 10.1016/j.chom.2013.04.009.

Miller, S.K., Good, R.T., Drew, D.R., Delorenzi, M., Sanders, P.R., Hodder, A.N., Speed, T.P., Cowman, A.F., de Koning-Ward, T.F., and Crabb, B.S. (2002). A subset of Plasmodium falciparum SERA genes are expressed and appear to play an important role in the erythrocytic cycle. J Biol Chem 277, 47524–47532. 10.1074/jbc.M206974200.

Moon, R.W., Hall, J., Rangkuti, F., Ho, Y.S., Almond, N., Mitchell, G.H., Pain, A., Holder, A.A., and Blackman, M.J. (2013). Adaptation of the genetically tractable malaria pathogen Plasmodium knowlesi to continuous culture in human erythrocytes. Proc Natl Acad Sci U S A 110, 531–536. 10.1073/pnas.1216457110.

Moreau, D., Vacca, F., Vossio, S., Scott, C., Colaco, A., Paz Montoya, J., Ferguson, C., Damme, M., Moniatte, M., Parton, R.G., et al. (2019). Drug-induced increase in lysobisphosphatidic acid reduces the cholesterol overload in Niemann-Pick type C cells and mice. EMBO Rep 20, e47055. 10.15252/embr.201847055.

Nasamu, A.S., Glushakova, S., Russo, I., Vaupel, B., Oksman, A., Kim, A.S., Fremont, D.H., Tolia, N., Beck, J.R., Meyers, M.J., et al. (2017). Plasmepsins IX and X are essential and druggable mediators of malaria parasite egress and invasion. Science 358, 518–522. 10.1126/science.aan1478.

Perrin, A.J., Collins, C.R., Russell, M.R.G., Collinson, L.M., Baker, D.A., and Blackman, M.J. (2018). The Actinomyosin Motor Drives Malaria Parasite Red Blood Cell Invasion but Not Egress. mBio 9. 10.1128/mBio.00905-18.

Pino, P., Caldelari, R., Mukherjee, B., Vahokoski, J., Klages, N., Maco, B., Collins, C.R., Blackman, M.J., Kursula, I., Heussler, V., et al. (2017). A multistage antimalarial targets the plasmepsins IX and X essential for invasion and egress. Science 358, 522–528. 10.1126/science.aaf8675.

Pszenny, V., Ehrenman, K., Romano, J.D., Kennard, A., Schultz, A., Roos, D.S., Grigg, M.E., Carruthers, V.B., and Coppens, I. (2016). A Lipolytic Lecithin:Cholesterol Acyltransferase Secreted by Toxoplasma Facilitates Parasite Replication and Egress. J Biol Chem 291, 3725–3746. 10.1074/jbc.M115.671974.

Reed, M.B., Caruana, S.R., Batchelor, A.H., Thompson, J.K., Crabb, B.S., and Cowman, A.F. (2000). Targeted disruption of an erythrocyte binding antigen in Plasmodium falciparum is associated with a switch toward a sialic acid-independent pathway of invasion. Proc Natl Acad Sci U S A 97, 7509–7514. 10.1073/pnas.97.13.7509.

Risco-Castillo, V., Topcu, S., Marinach, C., Manzoni, G., Bigorgne, A.E., Briquet, S., Baudin, X., Lebrun, M., Dubremetz, J.F., and Silvie, O. (2015). Malaria Sporozoites Traverse Host Cells within Transient Vacuoles. Cell Host Microbe 18, 593–603. 10.1016/j.chom.2015.10.006.

Ruecker, A., Shea, M., Hackett, F., Suarez, C., Hirst, E.M., Milutinovic, K., Withers-Martinez, C., and Blackman, M.J. (2012). Proteolytic activation of the essential parasitophorous vacuole cysteine protease SERA6 accompanies malaria parasite egress from its host erythrocyte. J Biol Chem 287, 37949–37963. 10.1074/jbc.M112.400820.

Schneider, C.A., Rasband, W.S., and Eliceiri, K.W. (2012). NIH Image to ImageJ: 25 years of image analysis. Nat Methods 9, 671–675. 10.1038/nmeth.2089.

Schultz, A.J., and Carruthers, V.B. (2018). Toxoplasma gondii LCAT Primarily Contributes to Tachyzoite Egress. mSphere 3. 10.1128/mSphereDirect.00073-18.

Silmon de Monerri, N.C., Flynn, H.R., Campos, M.G., Hackett, F., Koussis, K., Withers-Martinez, C., Skehel, J.M., and Blackman, M.J. (2011). Global identification of multiple substrates for Plasmodium falciparum SUB1, an essential malarial processing protease. Infect Immun 79, 1086–1097. 10.1128/IAI.00902-10.

Singh, P., Alaganan, A., More, K.R., Lorthiois, A., Thiberge, S., Gorgette, O., Guillotte Blisnick, M., Guglielmini, J., Aguilera, S.S., Touqui, L., et al. (2019). Role of a patatin-like phospholipase in Plasmodium falciparum gametogenesis and malaria transmission. Proc Natl Acad Sci U S A 116, 17498–17508. 10.1073/pnas.1900266116.

Thomas, J.A., Collins, C.R., Das, S., Hackett, F., Graindorge, A., Bell, D., Deu, E., and Blackman, M.J. (2016). Development and Application of a Simple Plaque Assay for the Human Malaria Parasite Plasmodium falciparum. PLoS One 11, e0157873. 10.1371/journal.pone.0157873.

Thomas, J.A., Tan, M.S.Y., Bisson, C., Borg, A., Umrekar, T.R., Hackett, F., Hale, V.L., Vizcay-Barrena, G., Fleck, R.A., Snijders, A.P., et al. (2018). A protease cascade regulates release of the human malaria parasite Plasmodium falciparum from host red blood cells. Nat Microbiol 3, 447–455. 10.1038/s41564-018-0111-0.

Thompson, J.K., Triglia, T., Reed, M.B., and Cowman, A.F. (2001). A novel ligand from Plasmodium falciparum that binds to a sialic acid-containing receptor on the surface of human erythrocytes. Mol Microbiol 41, 47–58. 10.1046/j.1365-2958.2001.02484.x.

Trager, W., and Jensen, J.B. (1976). Human malaria parasites in continuous culture. Science 193, 673–675. 10.1126/science.781840.

Viswanathan, S., Williams, M.E., Bloss, E.B., Stasevich, T.J., Speer, C.M., Nern, A., Pfeiffer, B.D., Hooks, B.M., Li, W.P., English, B.P., et al. (2015). High-performance probes for light and electron microscopy. Nat Methods 12, 568–576. 10.1038/nmeth.3365.

Wickham, M.E., Culvenor, J.G., and Cowman, A.F. (2003). Selective inhibition of a two-step egress of malaria parasites from the host erythrocyte. J Biol Chem 278, 37658–37663. 10.1074/jbc.M305252200.

Wirth, C.C., Glushakova, S., Scheuermayer, M., Repnik, U., Garg, S., Schaack, D., Kachman, M.M., Weissbach, T., Zimmerberg, J., Dandekar, T., et al. (2014). Perforin-like protein PPLP2 permeabilizes the red blood cell membrane during egress of Plasmodium falciparum gametocytes. Cell Microbiol 16, 709–733. 10.1111/cmi.12288.

Yague, G., Segovia, M., and Valero-Guillen, P.L. (1997). Acyl phosphatidylglycerol: a major phospholipid of Corynebacterium amycolatum. FEMS Microbiol Lett 151, 125–130. 10.1111/j.1574-6968.1997.tb12559.x.

Yang, A.S.P., O’Neill, M.T., Jennison, C., Lopaticki, S., Allison, C.C., Armistead, J.S., Erickson, S.M., Rogers, K.L., Ellisdon, A.M., Whisstock, J.C., et al. (2017). Cell Traversal Activity Is Important for Plasmodium falciparum Liver Infection in Humanized Mice. Cell Rep 18, 3105–3116. 10.1016/j.celrep.2017.03.017.

Yeoh, S., O’Donnell, R.A., Koussis, K., Dluzewski, A.R., Ansell, K.H., Osborne, S.A., Hackett, F., Withers-Martinez, C., Mitchell, G.H., Bannister, L.H., et al. (2007). Subcellular discharge of a serine protease mediates release of invasive malaria parasites from host erythrocytes. Cell 131, 1072–1083. 10.1016/j.cell.2007.10.049.

Zhang, M., Wang, C., Otto, T.D., Oberstaller, J., Liao, X., Adapa, S.R., Udenze, K., Bronner, I.F., Casandra, D., Mayho, M., et al. (2018). Uncovering the essential genes of the human malaria parasite Plasmodium falciparum by saturation mutagenesis. Science 360. 10.1126/science.aap7847.

Zimmerberg, J., and Kozlov, M.M. (2006). How proteins produce cellular membrane curvature. Nat Rev Mol Cell Biol 7, 9–19. 10.1038/nrm1784.

