## Supplementary figures and images for "A malaria parasite phospholipase facilitates efficient asexual blood stage egress"

### Supplemental Figure S1

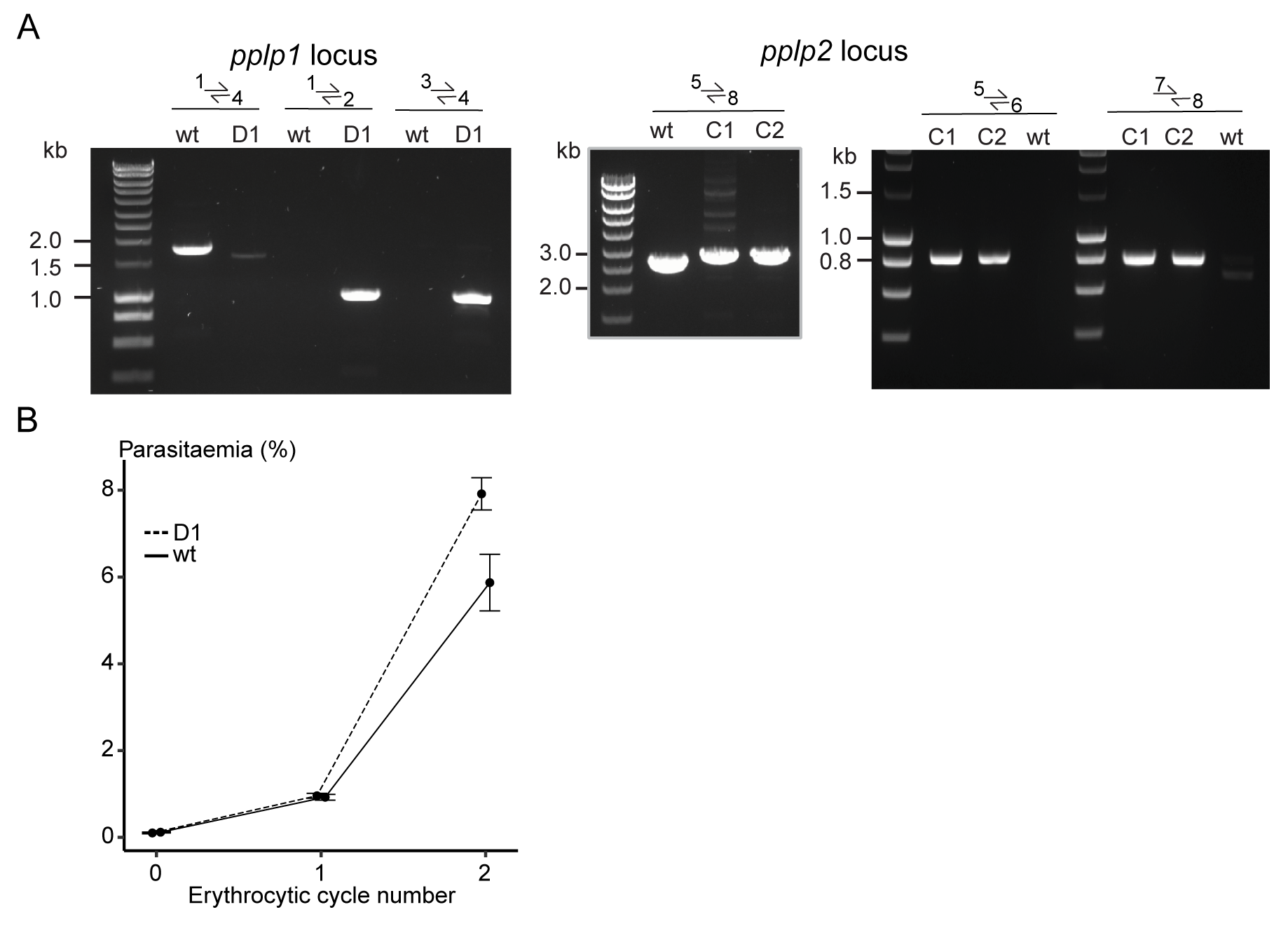

### Supplemental Figure S2

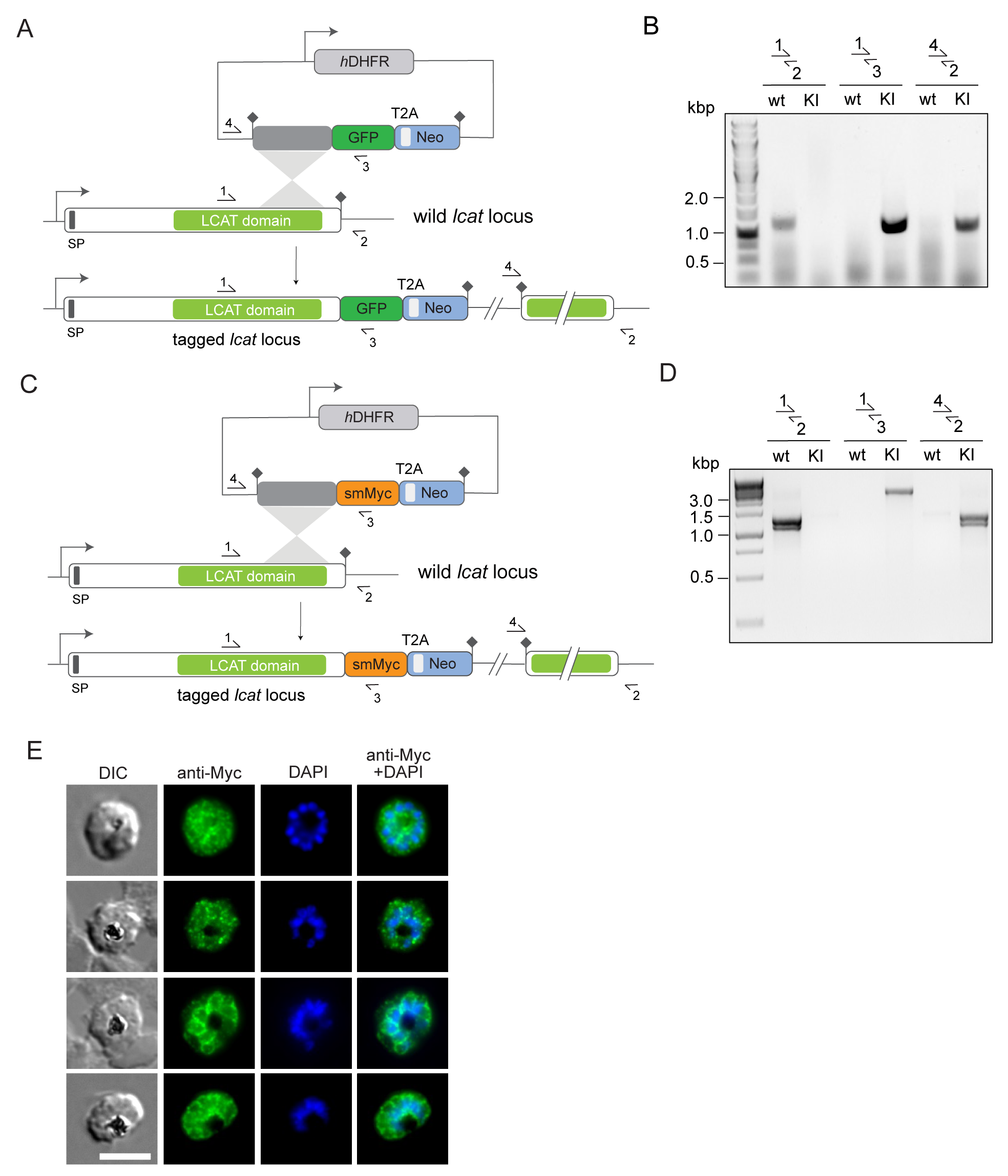

### Supplemental Figure S3

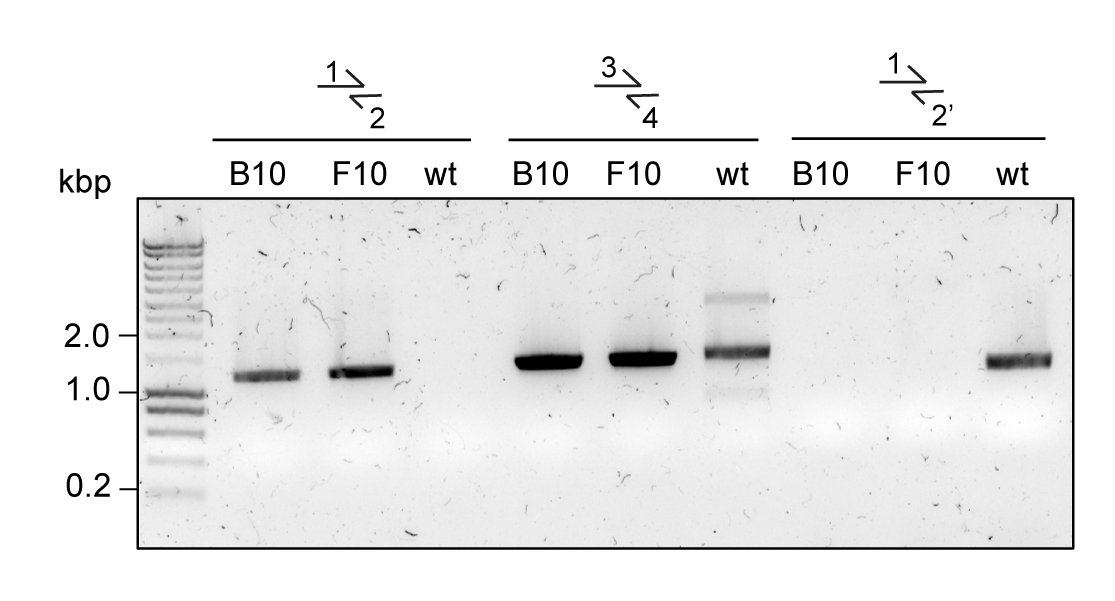

### Supplemental Figure S4

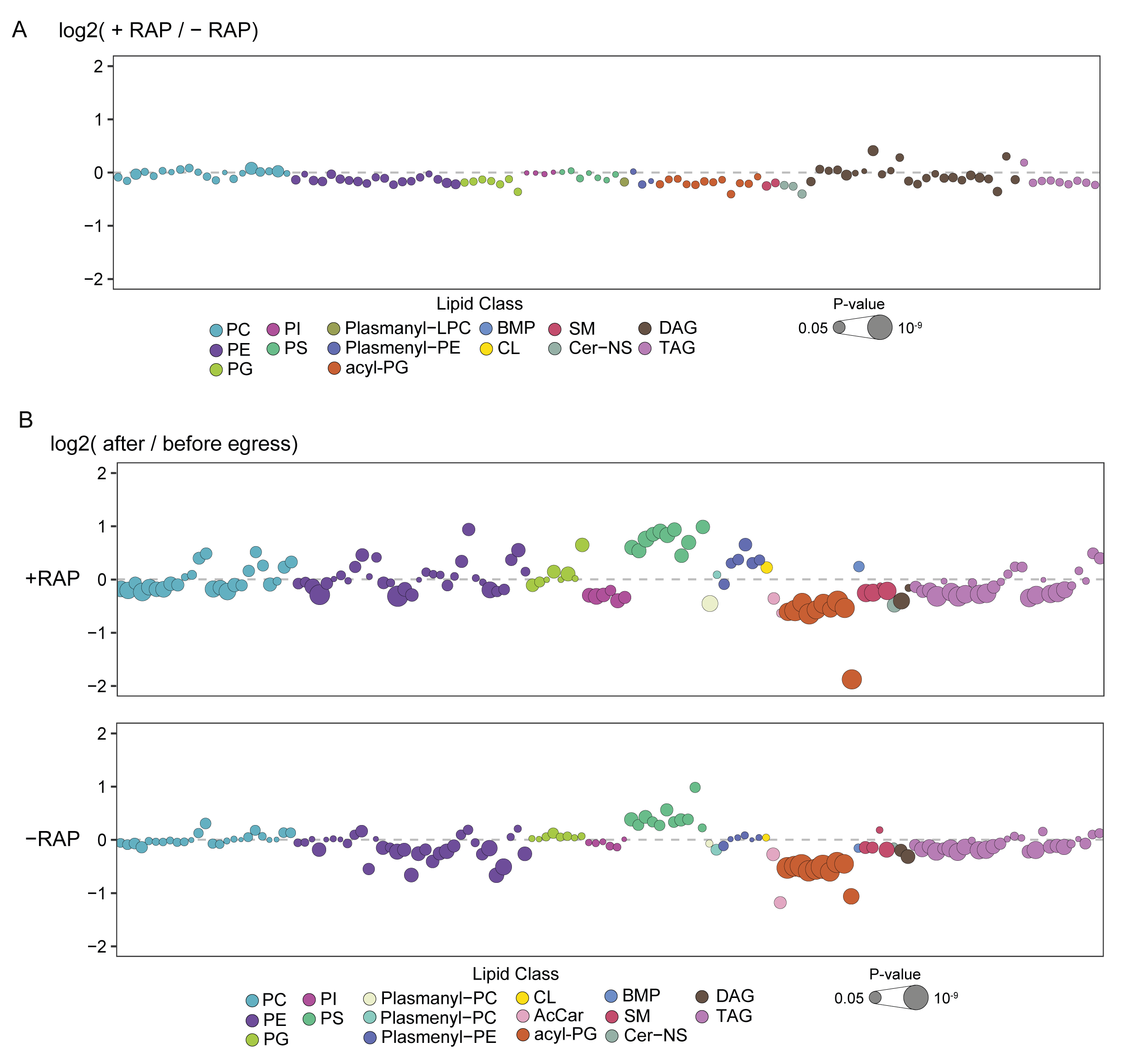

### Supplemental Figure S5

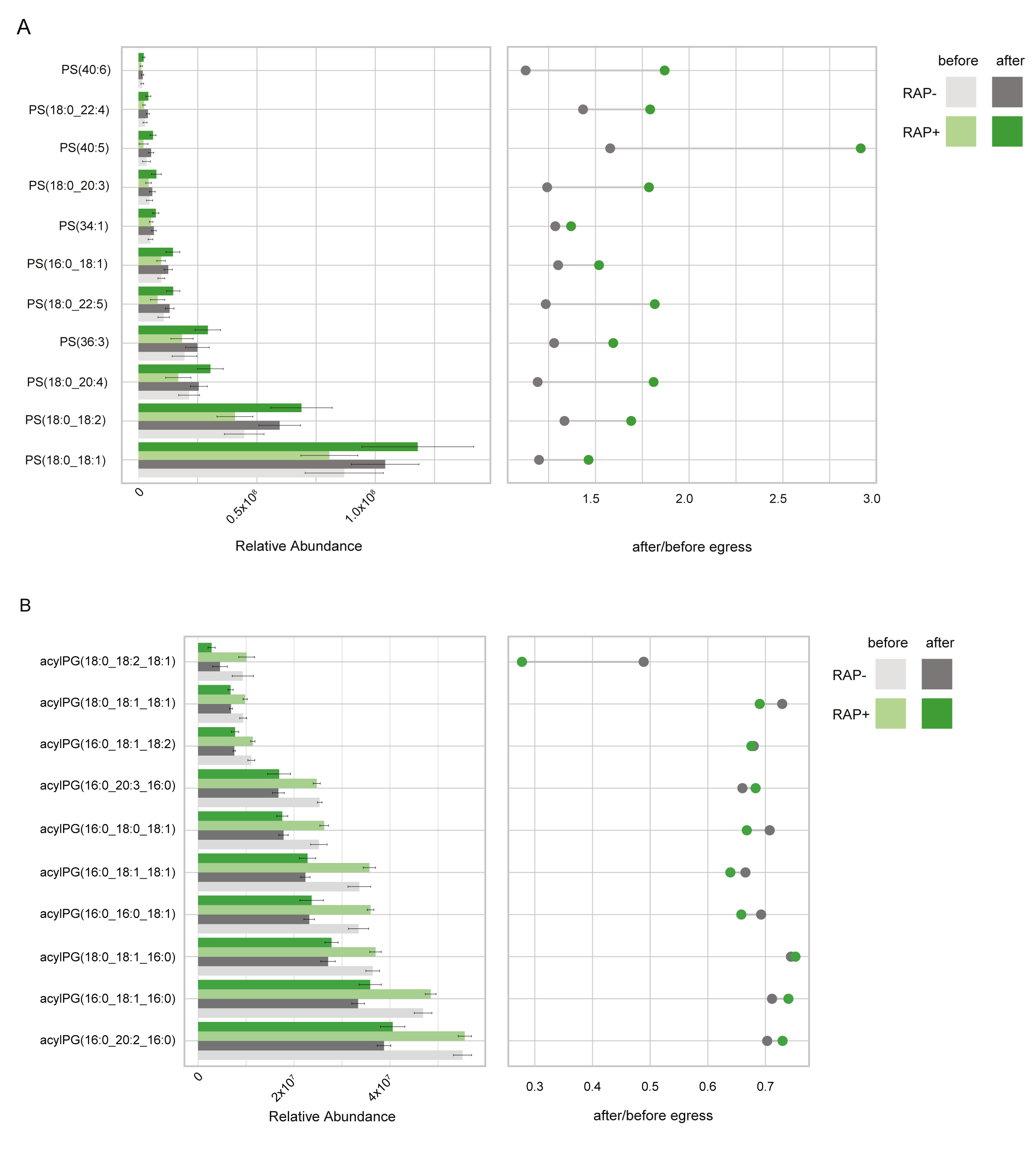
